# Genomic decoding of *Theobroma grandiflorum* (cupuassu) at chromosomal scale: Evolutionary insights for horticultural innovation

**DOI:** 10.1101/2023.12.23.573069

**Authors:** Rafael Moysés Alves, Vinicius A. C. de Abreu, Rafaely Pantoja Oliveira, João Victor dos Anjos Almeida, Mauro de Medeiros de Oliveira, Saura R. Silva, Alexandre R. Paschoal, Sintia S. de Almeida, Pedro A. F. de Souza, Jesus A. Ferro, Vitor F. O. Miranda, Antonio Figueira, Douglas S. Domingues, Alessandro M. Varani

**Affiliations:** Embrapa Amazônia Oriental, 66095-903 Belém, PA, Brazil; Laboratório de Bioinformática e Computação de Alto Desempenho (LaBioCad), Faculdade de Computação (FACOMP), Universidade Federal do Pará, 66075-110 Belém, PA, Brazil; Departamento de Biotecnologia Agropecuária e Ambiental, Universidade Estadual Paulista (UNESP), Faculdade de Ciências Agrárias e Veterinárias, 14884-900 Jaboticabal, SP, Brazil; Departamento de Biologia, Universidade Estadual Paulista (UNESP), Faculdade de Ciências Agrárias e Veterinárias, 14884-900 Jaboticabal, SP, Brazil; Departamento de Ciência da Computação (DACOM), Grupo de e Bioinformática e Reconhecimento de Padrões (bioinfo-cp), Universidade Tecnológica Federal do Paraná (UTFPR), 80230-901 Cornélio Procópio, PR, Brazil; Artificial Intelligence and Informatics, The Rosalind Franklin Institute, Didcot, UK; Centro de Energia Nuclear na Agricultura (CENA), Universidade de São Paulo, Piracicaba, SP, Brazil; Departamento de Genética, Universidade de São Paulo (USP), Escola Superior de Agricultura Luiz de Queiroz (ESALQ), Piracicaba, SP, Brazil

**Keywords:** Amazon basin, bioeconomy, fruit pulp and seed development, genome evolution, gene loss and retention, positive selection, plant secondary metabolites

## Abstract

**Background:** *Theobroma grandiflorum* (Malvaceae), known as cupuassu, is a tree indigenous to the Amazon Basin, valued for its large fruits and seed-pulp, contributing notably to the Amazonian bioeconomy. The seed-pulp is utilized in desserts and beverages, and its seed butter is used in cosmetics. Here, we present the sequenced telomere-to-telomere cupuassu genome, disclosing features of the genomic structure, evolution, and phylogenetic relationships within the Malvaceae.

**Results:** The cupuassu genome spans 423 Mb, encodes 31,381 genes distributed in the ten chromosomes, and it exhibits approximately 65% gene synteny with the *T. cacao* genome, reflecting a conserved evolutionary history, albeit punctuated with unique genomic variations. The main changes are pronounced by bursts of long-terminal repeats retrotransposons expansion at post-species divergence, retrocopied and singleton genes, and gene families displaying distinctive patterns of expansion and contraction. Furthermore, positively selected genes are evident, particularly among retained and dispersed, tandem and proximal duplicated genes associated to general fruit and seed traits and defense mechanisms, supporting the hypothesis of potential episodes of subfunctionalization and neofunctionalization following duplication, and impact from distinct domestication process. These genomic variations may underpin the differences observed in fruit and seed morphology, ripening, and disease resistance between cupuassu and the other Malvaceae species.

**Conclusions:** Sequencing the cupuassu genome offers a foundational resource for both breeding and conservation efforts, yielding insights into the evolution and diversity within the genus *Theobroma*.

**Core ideas:**

- Telomere-to-telomere sequencing of the *Theobroma grandiflorum* genome elucidates a 65% synteny with *T. cacao*.
- Retrotransposon expansion identified as a pivotal factor in post-divergence genomic evolution between *Theobroma* species.
- Comparative genomics has revealed genes associated with key agronomic traits, providing evolutionary insights.
- Positive selection pressure in retained duplicated genes implicated in adaptive functions and fruit-seed traits diversity.
- Cupuassu genome as a genetic resource for breeding and to boosts Brazillian Amazonian bioeconomy.

**AUTHOR SUMMARY:** Cupuassu, a fruit-bearing tree from the Amazon, is prized for its nutritious fruits and seeds, widely used in food and cosmetics. In this study, we sequenced the complete genome of cupuassu to understand its development, unique traits, and genetic relationship with cacao and other related species. The cupuassu genome shares a high similarity with cacao, but it also exhibits distinctive features. Notably, repetitive DNA elements have significantly influenced its genomic structure. Furthermore, specific genes responsible for its fruit and seed characteristics, as well as disease resistance, were identified. Overall, this research not only deepens our knowledge of cupuassu genetics but also illuminates broader aspects of plant evolution and diversity in the Amazon. It lays the groundwork for advanced breeding programs and promises to contribute significantly to the Amazonian bioeconomy. Ultimately, these findings have important implications for agriculture and conservation, highlighting the intricate processes of plant adaptation and evolution.

## Background

The genus *Theobroma* L. (Malvaceae) originated in the Neotropical regions, with the Amazon basin as its main ecosystem. Among the 22 *Theobroma* species [1,2], two species, *T. cacao* (cacao) and *T. grandiflorum* (cupuassu) are of significant economic importance. Both are diploid (2*n* = 2× = 20) presenting an average genome size around 450 Mb [3]. These species display distinct fruit and seed morphologies, which are likely the most valued parts by humans and other dispersers [4]. Cacao seeds are the main component for the chocolate and confectionery industries. In contrast, cupuassu seed pulp is used in desserts and beverages. Additionally, cupuassu seeds can be processed to create a butter highly prized in the cosmetic industry and ‘cupulate,’ a product akin to chocolate [5].

Cupuassu, domesticated from *T. subincanum* by Amazon indigenous populations approximately 5,000 to 8,000 years ago, has spread geographically mainly in the last two centuries [6]. In Brazil, cupuassu is especially important for small-scale farmers in agroforest systems in Pará, Amazonas, and Bahia, the leading states in its production [7]. In 2022, Brazilian cupuassu production reached about 28,800 tons of fresh seeds from 8,900 hectares, averaging 3.2 tonnes per hectare (State Secretariat for Agricultural Development. Agricultural Indicators. Belém, PA, Brazil, 2022).

Both cacao and cupuassu face substantial threats from various fungal and viral pathogens. Specifically, the witches’ broom disease (WBD) and frosty pod (FP) pose major challenges. Both diseases are caused by two basidiomycete species, *Moniliophthora perniciosa* and *M. roreri*, respectively. These diseases significantly reduce pod yield and the overall health of infected plants, resulting in substantial economic losses [8]. While breeding programs have developed resistant cacao and cupuassu genotypes [7,8], managing WBD and FP remains challenging [9,10], impacting the local producers and family farmers systems.

Numerous sequencing initiatives have been undertaken for cacao to provide insights into the genome biology, plant-pathogen interactions and to assist breeding over the past 15 years [11–16]. To date, 37 chromosome-scale *T. cacao* genomes are publicly accessible, encompassing a range of genotypes from widely cultivated to wild-collected accessions. Additionally, the genome sequence of *Herrania umbratica*, a sister genus to *Theobroma* known as ‘monkey cacao,’ which exhibits unique morphology [17], is also available.

In parallel, recent investigations have delved into the genomic architecture of *T. grandiflorum*, ranging from the first genetic map [18], the chloroplast and mitochondrial genomes [19,20], and comparative transcriptomics [10,21]. These latter studies shed light on the interaction between cupuassu and *M. perniciosa*, setting the groundwork for breeding programs and transgenic approaches. However, limited genomic data for *T. grandiflorum* persists, leaving gaps in understanding its genome evolution, biology and potential comparison with *T. cacao*, a key crop in the genus.

In this study, we present a detailed analysis of the *T. grandiflorum* genome, assembling a high-quality telomere-to-telomere (T2T) chromosome-scale genome. Our comparative genomic approach reveals important genomic features, distinguishing it from closely related species like *T. cacao* and *H. umbratica*. These insights provide critical targets for breeding and of significant importance for evolutionary biology, biotechnology, conservation and horticulture research.

## Methods

### Plant sampling, DNA and RNA extraction, and sequencing

Leaf samples of the cupuassu clone 1074, susceptible to WBD [18], were collected at the ‘Embrapa Amazônia Oriental’ collection in Belém, PA, Brazil (1.4359° S, 48.4495° W), and cataloged at the Herbarium JABU, Universidade Estadual Paulista, Jaboticabal campus (Voucher JABU1370). The samples underwent a 24-h dark incubation, flash freezing in liquid nitrogen, and were transported to the Arizona Genomics Institute (Tucson, USA) for analysis. High molecular weight (HMW) DNA was extracted using a modified CTAB protocol [22], assessed for integrity and concentration via Qubit dsDNA High-Sensitivity Assay (Thermo Fisher Scientific, Waltham, MA, USA) and NanoDrop NP-1000 (NanoDrop Technologies, Wilmington, DE, USA). DNA quality and size were confirmed with Femto Pulse and pulse-field gel electrophoresis (Femto Pulse System, Agilent Technologies, Inc, Santa Clara, CA, USA). The DNA was sheared to 10–30 Kb using a Covaris g-TUBE (Covaris, Inc, Woburn, MA, USA), purified, and sequenced on a PacBio Sequel IIe platform (PacBio, Menlo Park, CA, USA). GenomeScope 2.0 [23] and KMC v3.2.1 [24] were employed for genome profiling.

Total RNA was extracted using the PureLink Plant RNA Reagent (Thermo Fisher Scientific Inc) and Takara NucleoSpin® RNA Clean-up(Takara Bio Inc, Kusatsu, Shiga, Japan). RNA integrity was confirmed by a 2100 Bioanalyzer (Agilent Technologies, Santa Clara, CA, USA), and only samples with an RNA Integrity Number above 7 proceeded to sequencing. IsoSeq library preparation and sequencing were performed on a PacBio Sequel IIe, while Illumina sequencing was conducted on a HiSeq 2000 platform (Illumina, Inc, San Diego, CA, USA) at NGS Soluções Genômicas, Brazil.

For HiC library preparation and sequencing, samples were processed at Novogene Bioinformatics Technology (Beijing, China) using the Proximo™ Hi-C Kit (Seattle, WA, USA). The quality control was conducted using Phase Genomics’ hic_qc scripts (https://github.com/phasegenomics/hic_qc).

### Genome assembly and quality evaluation

PacBio HiFi reads were assembled employing Hifiasm v0.19.3-r572 [25] with default parameters. Contaminants were removed using kraken2 [26] and “extract_kraken_reads.py” (https://github.com/jenniferlu717/KrakenTools), with the PlusPFP index database (https://benlangmead.github.io/aws-indexes/k2). The primary assembly was indexed with BWA v0.7.17-r1188 [27], and *Dpn*II restriction sites were created using the Juicer pipeline v1.6 [28]. Genome scaffolding and chromosomal reconstruction were achieved using 3D-DNA v180419 [29], and manually corrected with Juicebox Assembly Tools v3.1.4 [28]. The final chromosome-level assembly was refined using the “run-ASM-pipeline-post-review.sh” script from 3D-DNA and “close_scaffold_gaps.sh” from the MaSuRCA assembler package [30].

For the *H. umbratica* accession Fairchild (BioProject: PRJNA383741), re-assembly used the MaSuRCA hybrid approach with PacBio CLR and Illumina reads. Genome scaffolding for this genotype employed the Arima Genomics’ mapping pipeline (https://github.com/ArimaGenomics/mapping_pipeline) and YaHS [31].

The assembled genome quality and completeness were validated using Merqury v1.3 [32], LTR Assembly Index (LAI) [33], and BUSCO (Benchmarking Universal Single-Copy Orthologs) v5.4.5 against the embryophyta_odb10 database [34,35].

### Transcriptome and IsoSeq Assembly

IsoSeq transcripts fasta file was generated using SMRT Link 12.0 (PacBio) with default parameters. *De novo* assembly of RNAseq short-reads and HiFi reads were performed using Trinity pipeline v2.14.0 [36]. For genome-guided transcriptome assembly, the short-reads and HiFi reads were separately aligned to the chromosome-scale genome using histat2 v2.2.1 [37] and minimap2 v2.24-r1122 [38], respectively. The aligned BAM files from both read types were then merged using StringTie2 v2.2.1 [39] to produce a GTF file, which was utilized in the genome annotation process.

The PASA v2.5.3 pipeline [40], integrating IsoSeq fasta, *de novo*, and genome-guided assemblies with StringTie2, along with TransDecoder v5.7.0 (Haas, BJ. https://github.com/TransDecoder/TransDecoder) were employed to create a comprehensive transcriptome database and annotate transcript structures. This methodology was applied to both *T. cacao* v2 (Belizian Criollo B97-61/B2 cultivar) [12] and *H. umbratica* (Fairchild) transcriptomes, using public short-reads from the GenBank Sequence Read Archive (Table S1).

The completeness of the assembled transcriptome was assessed using BUSCO v5.4.5 against the embryophyta_odb10 database in transcriptome mode.

### Genome Annotation and Comparative Analyses

The genome annotation was carried out in two phases, following best practices in plant genome annotation [41]. Detailed procedures are provided in Supplementary Information 1.

The genome map was created using shinyCircos-V2.0 [42]. Whole-genome duplication (WGD) and positive selection analyses followed established methods [43]. Macrosynteny and microsynteny were analyzed using MCScanX [44], SynVisio [45], and the Python version of Mcscan [46], with synteny percentages computed using custom Python scripts based on MCscan outputs.

Chromosome plots were generated with the jcvi miscellaneous plotting tool (https://github.com/tanghaibao/jcvi/wiki/Miscellaneous-plotting) and the MG2C tool [47]. Transposable elements (TE) distribution relative to genes was determined using TE_Density [48]. The TE distribution plot was generated with RAWgraphs v2.0 [49]. Orthologous gene clusters (gene families) were identified using OrthoVenn3 [50] and the OrthoFinder2 algorithm [51] with diamond in super-sensitive mode [52]. Gene family evolution was analyzed using CAFE 5 [53]. For comparative purposes and to root the phylogenetic tree, the cotton D genome (*Gossypium raimondii*) v. 2.1 [54] and *Arabidopsis thaliana* (version Araport11) [55] were employed.

Gene Ontology (GO) enrichment analyses were performed with GOATOOLS [56], considering only results with a p-value below 0.05 after false discovery rate correction with Benjamini/Hochberg significance test. Targeted comparative analyses focused on genes and functions previously related to seed traits and fruit characteristics, such as aroma, quality, maturation, and flavor, incorporating components like purine alkaloids, flavonoids, terpenoids, and fatty acids [11,57]. This was supplemented by literature and Gene Ontology searches through the QuickGO platform [58].

### Data availability

The *T. grandiflroum* sample (GenBank BioSample SAMN37717187) was included at National Genetic Heritage and Associated Traditional Knowledge Management System (SisGen) under the accession #A2A72C6. The complete genome was deposited at GenBank, BioProject PRJNA691024 the raw reads are available at GenBank Sequence Read Archive (SRA) under the accession numbers: SRR26316970 and SRR26316971. The genome sequence, gene models and functional annotation files (GFF3s and FASTAs) are also available at our genome browser web-service: https://plantgenomics.ncc.unesp.br/gen.php?id=Theo.

## Results and Discussion

### High-Resolution Chromosome-Level Genome Assembly of *T. grandiflorum*

The *T. grandiflorum* assembled genome contains 10 chromosomes, each ranging from 28 to 53 Mb and cumulatively spanning 423,916,809 bp, with an average read coverage of 40x (Figure 1A and B, Table 1, and Table S2). This assembly represents approximately 94% of the haploid genome size estimated by flow cytometry [3]. Telomeric repeats AAACCCT/AGGGTTT were identified in the chromosomes extremities indicating that T2T assembly was achieved. The average GC content of the cupuassu genome is 34.01%, comparable to *H. umbratica* (33.76%) and to *T. cacao* (32.14%). The chromosome-level assembly displays a consensus quality of 68, an error rate of 1.61e^−7^, k-mer completeness score of 88%, LAI of 15.6, a BUSCO score of 98.4%, and a k-mer heterozygosity rate of 0.61%.

**Figure 1.**
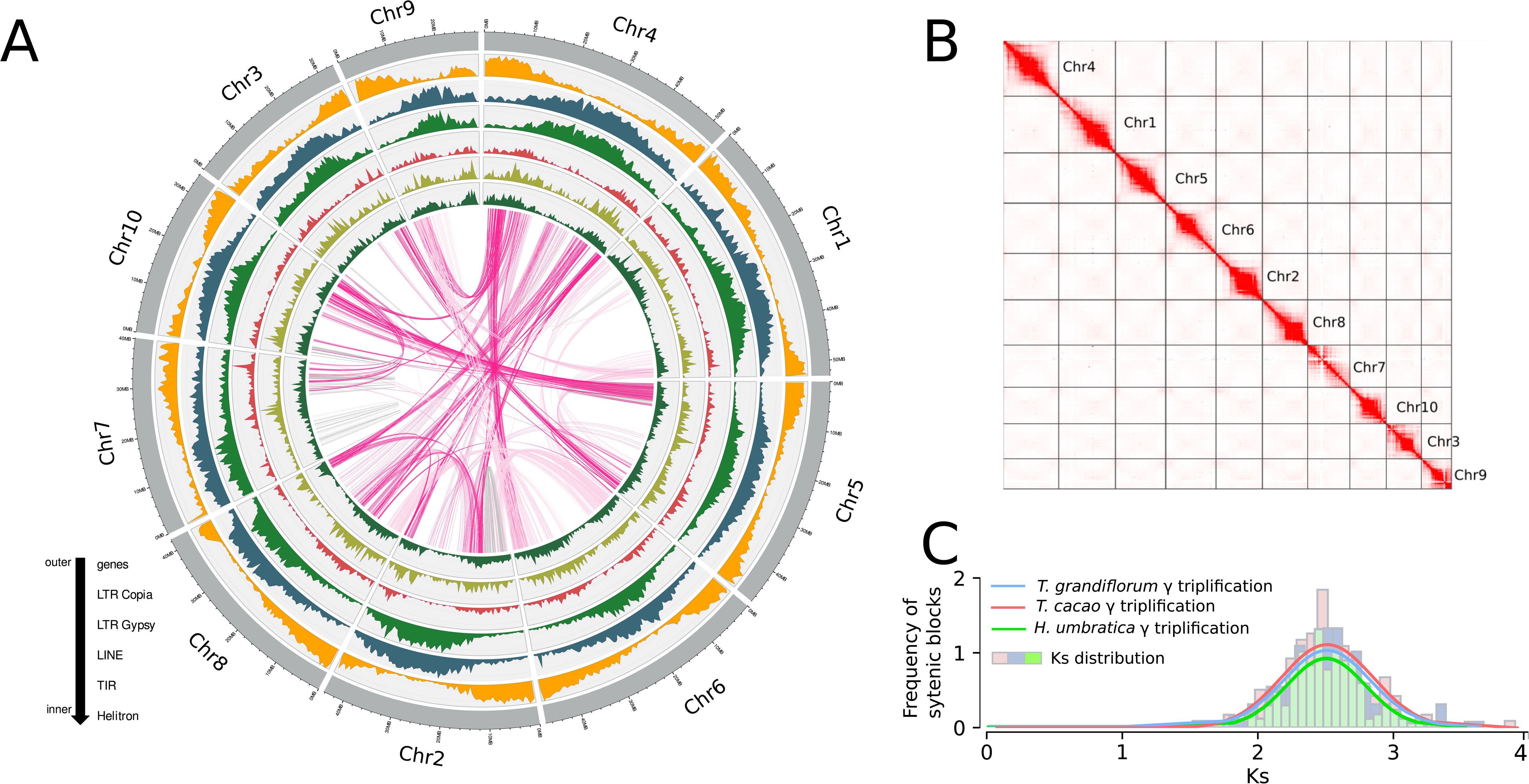
**A.** Depiction of the genomic landscape of *Theobroma grandiflorum*, illustrating gene and TE density across the ten chromosomes. **B.** High-throughput chromosome conformation capture (Hi-C) contact map revealing the assembled chromosomes of *T. grandiflorum*. **C.** Whole genome duplication analyses indicating the shared whole genome triplication among *T. grandiflorum*, *T. cacao*, and *H. umbratica*, and confirming the absence of additional WGD events in these species.

**Table 1.** *Theobroma grandiflorum* genome and annotation features.

| Features | *T. grandiflorum* |
| --- | --- |
| -Genome length | 423,916,809 bp |
| -GC% | 34.01 |
| -Number of genes | 31,381 |
| -- Number of CDSs | 46,671 |
| ---- complete CDS | 46,625 |
| ---- start, no stop CDS | 8 |
| ---- stop, no start CDS | 22 |
| ---- no stop, no start CDS | 16 |
| -mean gene length | 3,374 bp |
| -mean CDS length | 1,331 bp |
| -mean exons per gene | 6 |
| -mean introns per gene | 5 |
| -tRNAs | 446 |
| -snRNAs | 976 |
| -miRNAs | 109 |
| % of genome covered by genes | 25% |
| % of genome covered by CDS | 14.70% |
| % of genome covered by TEs | 53.35% |
| ---Class I Elements | 43.86% |
| LTR Gypsy | 13.18% |
| LTR Copia | 18.31% |
| LTR non-autonomous | 12.37% |
| non-LTR | 0.58% |
| ---Class II Elements | 2.36% |
| TIRs | 1.21% |
| Helitron | 1.15% |
| ---Other repeats | 7.13% |

### Structural Annotation and Spatial Gene Arrangement

A total of 31,381 protein-coding genes, corresponding to up 25% of the entire cupuassu genome, were identified (Table 1 and Table S3). The structural gene annotation achieved a BUSCO completeness of 99.8%. Through RNAseq and IsoSeq reads mapping, 46,625 complete coding sequence (CDS) were determined, confirming the functional isoforms in the gene models. The average gene length was 3,374 bp and CDS length 1,331 bp with 6 exons, values similar to *T. cacao* [11]. Furthermore, their distribution is evenly spread across the ten chromosomes.

The gene spatial arrangement and distribution in *T. grandiflorum* and *T. cacao* genomes show a similar pattern accordingly to their closely related evolutionary ties. This pattern includes genes from various duplications (whole-genome, tandem, proximal, transposed, dispersed) (Table S3). Analysis of Ks values and WGD-gene pairs distribution within syntenic blocks, using Gaussian mixture models, revealed a distinct Ks peak at 2.5. This finding corroborates the core eudicot γ whole-genome triplication event (Figure 1 C).

The cupuassu genome contains 402 genes that have originated through RNA-mediated duplication, referred to as retrocopies, comprising 197 chimeric genes, 37 pseudogenes, and 168 retrogenes. A comparative analysis of these retrocopies with *T. cacao* and *H. umbratica* highlighted unique retrocopies in each species: 67 in *T. grandiflorum*, 50 in *T. cacao*, and 34 in *H. umbratica* (Table S4). Interestingly, some of the unique retrocopies are linked to potential fruit and seed quality traits and plant development. For instance, a number of exclusive retrocopies in these species are related to serine/threonine-protein kinase, which is important for signal transduction and plays relevant roles in pathogen defense and fruit abscission [59]. Furthermore, retrocopies associated with chalcone metabolism in *T. grandiflorum* (TgrandC1074G00000001563) and embryo sac development in *T. cacao* (Tcacao-CriolloG00000031869) were also identified. Additionally, unique retrocopied transcription factors were noted, such as an auxin response factor in *T. grandiflorum* (TgrandC1074G00000000856) and WER-like transcription factors in *T. cacao* (Tcacao-CriolloG00000008811). *H. umbratica* unique retrocopies include genes linked to a caffeic acid 3-O-methyltransferase-like activity (HumbraticaG00000009034) and polygalacturonase (HumbraticaG00000026833), potentially influencing fruit traits.

In *T. cacao*, ncRNAs have been proposed as primary regulators of gene expression [11]. In cupuassu, our annotation identified 1,178 long non-coding RNAs (lncRNAs), 1,058 small nucleolar RNAs (snoRNAs), 446 transfer RNAs (tRNAs), 126 microRNAs (miRNAs), 48 small nuclear RNAs (snRNAs), and 17 small RNAs (sRNAs). Moreover, the primary sites for 5S and 45S ribosomal DNA (rDNA) were mapped to chromosomes 2 and 7, respectively, corroborating previous rDNA localization using fluorescent *in situ* hybridization [60]. Overall, ncRNAs are relatively evenly distributed across the chromosomes. Notably, chromosome 7 has the lowest counts of tRNAs, miRNAs, and sRNAs (Figure S1 and Table S5).

### TE Distribution and Impact in the Cupuassu Genome Architecture and Function

TE and repetitive elements constitute roughly 53% of the *T. grandiflorum* genome. The most abundant TE were LTR *Copia*, LTR *Gypsy*, and the non-autonomous LARD elements (Figure 2 A and Table S6). Notably, LTR *Copia* SIRE and LTR *Gypsy* Tekay were the most prevalent lineage accounting for up to 49 and 36Mb of the genome (Figure 2B). Evolutionarily, LTR *Copia* elements had two significant peaks of expansion at 0.3 and 1.8 million years ago (mya), whereas the LTR *Gypsy* elements showed a single peak around 0.3 mya (Figure S2). In general, the LTR insertion ages shows similar pattern in several plant families, like Fabaceae, Solanaceae, Poaceae, Funariceae, Salicaceae, Musaceae, Selaginellaceae, and Brassicaceae [61].

**Figure 2.**
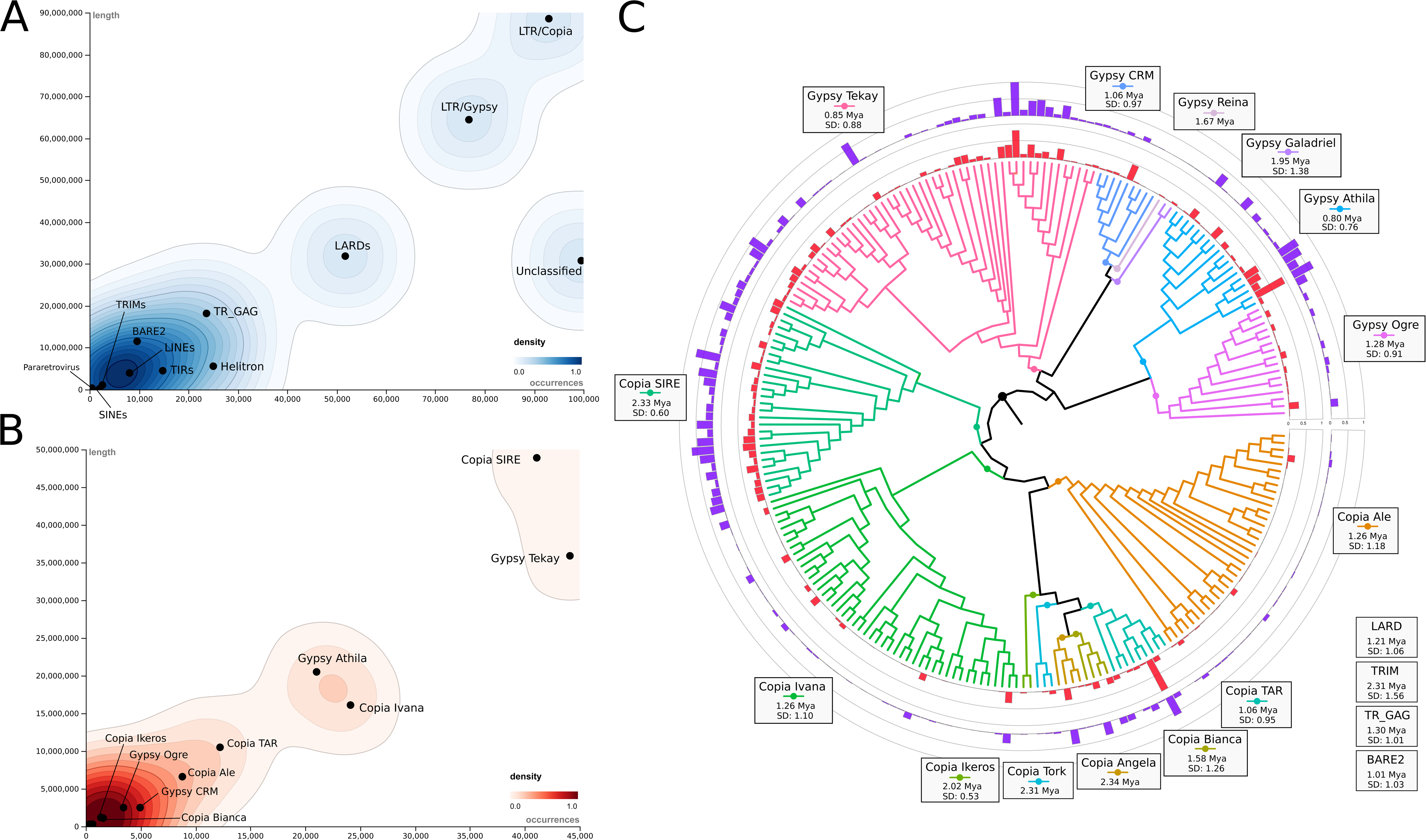
Transposable Elements Distribution in *Theobroma grandiflorum.* **A.** Distribution of autonomous and non-autonomous TE from Class I and Class II. **B.** Distribution of all evolutionary lineages of LTR elements. **C.** Phylogenetic analysis and distribution of each full-length LTR element identified in *Theobroma grandiflorum*.

While the LTR *Copia* SIRE and LTR *Gypsy* Tekay elements are notably abundant in *T. grandiflorum*, they display unique expansion pattern and ages (Figure 2 C). Almost all members of *Copia SIRE* exhibit expansion, whereas only a subset of *Gypsy Tekay* elements show similar expansive trend. In contrast, certain LTR lineages, particularly *Copia TAR* and *Gypsy Athila*, have undergone significant proliferative events, marking their distinctive expansion. Interestingly, despite the high membership of *Copia Ivana*, *Ale*, and *Gypsy Ogre*, these lineages exhibit limited proliferation. In contrast, the Class II elements were less prominent as observed in other plant genomes, including *T. cacao* [11,62]. For instance, the MuDR/Mutator lineage is the most abundant, covering 881 Kb (0.27%) of the cupuassu genome.

The distribution of TE across cupuassu chromosomes is uniform among all TE classes and lineages (Table S5). The density of TEs around gene regions reflects their overall abundance in the genome, with LTR *Copia*, LTR *Gypsy*, and LARDs being concentrated near genes, typically located about 1.5 Kb both upstream and downstream (Figure S3). This distribution pattern supports the idea that TEs are advantageously placed, rather than randomly, possibly impacting gene expression regulation and influencing regulatory networks [63].

### *Theobroma grandiflorum* exhibits elevated syntenic relationships with cacao, and *H. umbratica*

At the macrosyntenic level, both *Theobroma* species exhibit significant genomic conservation, suggesting minimal rearrangements (Figure 3 A), an observation that aligns with a previously published high-density cupuassu genetic map [18]. Notable variation occurs primarily within the pericentromeric and predicted centromeric regions, characterized by an elevated TE density, and other TE dense regions (Figure 3 B). This pattern is consistent with what is commonly found in plant genomes and it has been previously observed in the cacao genome [11].

**Figure 3:**
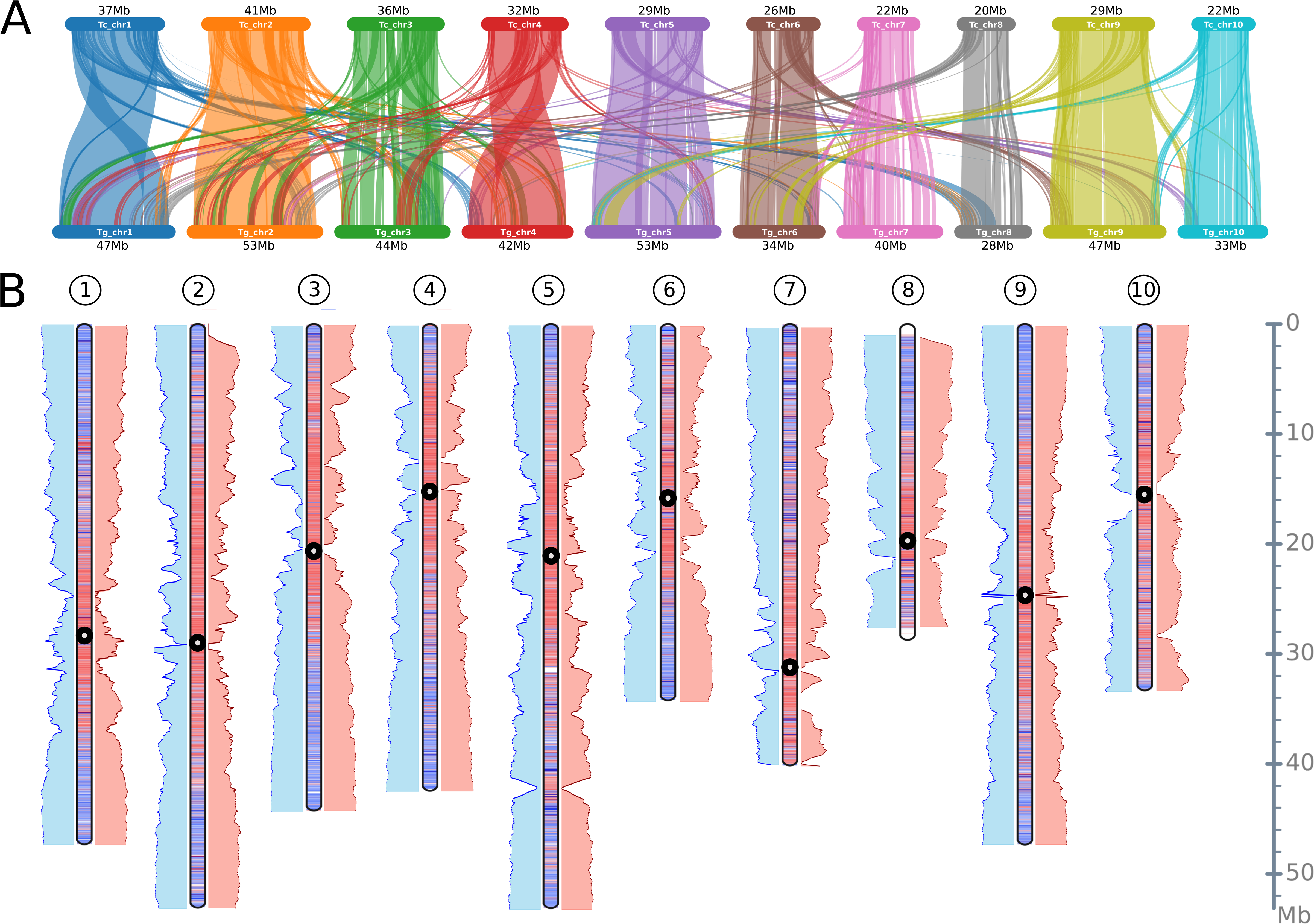
Comparative Genomic Analysis of *Theobroma grandiflorum* with *T. cacao* and *Herrania umbratica.* **A.** Macrosyntenic patterns between *Theobroma grandiflorum* and *T. cacao*, revealing conserved genome structures. **B.** Comparative idiogram map between *Theobroma grandiflorum* and *T. cacao*, and between *T. grandiflorum* and *Herrania umbratica*. The idiograms illustrate gene-rich regions (blue), TE-rich regions (red), and potential location of centromeres (black circles). Blue bars on the left of each idiogram represent microsynteny between *T. grandiflorum* and *T. cacao*, while red bars on the right indicate microsynteny between *T. grandiflorum* and *H. umbratica*.

A closer inspection at the microsyntenic level among *T. grandiflorum*, *T. cacao*, and *H. umbratica* reveals a marked gene synteny and collinearity, especially within the subtelomeric regions (Figure 3B and Figure S4). The three species preserves at least 65% (Table S5). Interestingly, transposed gene pairs between these species are comparatively infrequent (around 7% on average).

### Microsyntenic Insights into the Self-Incompatibility Loci of *Theobroma* and *Herrania*

Previous research identified two self-incompatibility loci in cacao, CH1 and CH4, with CH4 primarily linked to fruit drop [64]. Microsyntenic comparison of these loci in *T. grandiflorum* and *H. umbratica* revealed distinct patterns (Figure 4 A and B). CH1 is highly conserved across the three species, except for a missing COMPASS-like H3K4 histone methylase gene in *H. umbratica*, crucial in cellular networks [65]. CH4, however, varies significantly; *T. grandiflorum* and *H. umbratica* sequences are conserved, but the one in*T. cacao* contains two additional, truncated GEX1 gene copies (Figure S5), influencing gametophyte and embryo development and possibly affecting fruit setting and late incompatibility in *T. cacao* [64,66]. The CH4 locus in cacao also feature many TE remnants and a complete LTR-RT from the Copia/Tork lineage near the GEX1 gene.

**Figure 4:**
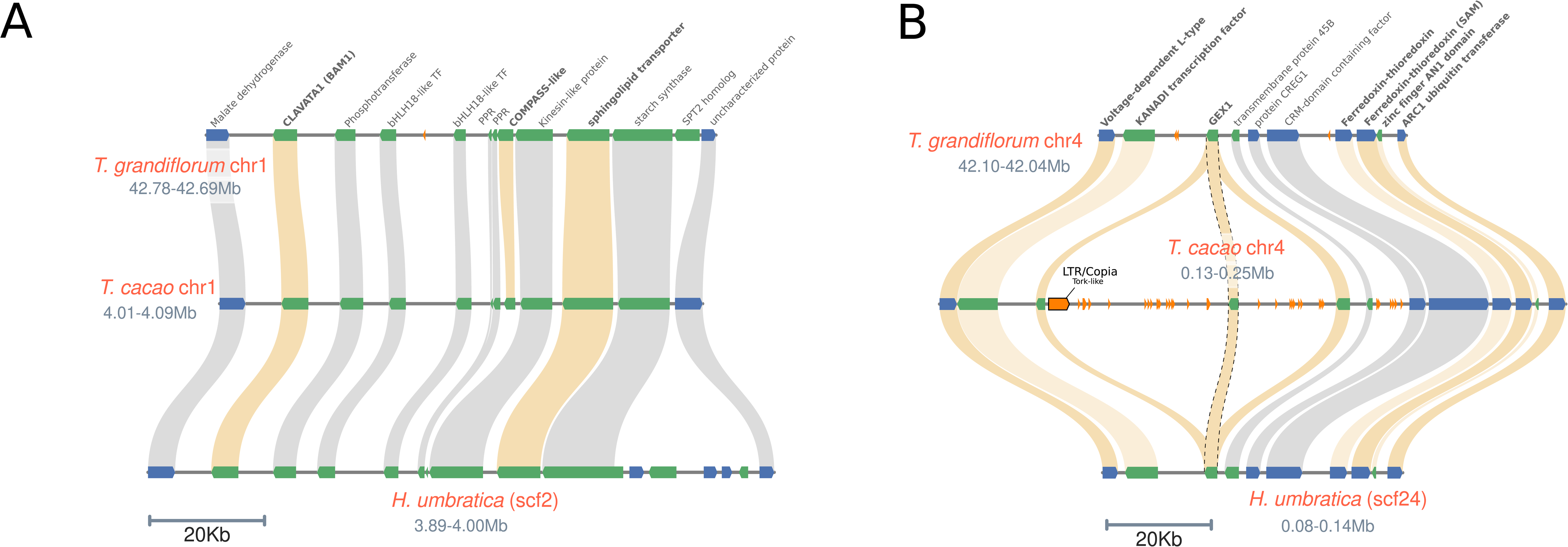
Microsyntenic Analysis of the Self-Incompatibility Loci (CH1 and CH4) in Theobroma and *Herrania.* **A.** CH1 loci. **B.** CH4 loci. Genes marked in bold are considered central to self-incompatibility reactions, as previously described [64]. The GEX1 locus, containing the complete and homologous genes, is marked with dotted lines.

Cupuassu and cacao are notably different for fruit abscission. Cupuassu fruits fall naturally when ripe, whereas cacao fruits need to be harvested from the tree [67,68]. We speculate that the multiple copies of the cacao *GEX1* gene, including the two truncated ones, in conjunction with the proximity of TE at the CH4 loci, could either affect *GEX1* expression or produce non-functional *GEX1* proteins. This potential influence may be linked to the fruit abscission phenotype in cacao, though it necessitates further empirical investigation for confirmation.

### Comparative Analyses Reveal Exclusive Cupuassu Genes and Distinct Patterns of Gene Family Expansion and Contraction associated with fruit quality traits and defense mechanisms

A total of 282 exclusive gene families and 1,160 singletons were identified in *T. grandiflorum* (Figure 5 A), whereas 730 gene families are shared between *T. grandiflorum* and *T. cacao*, and 297 gene families are shared between *T. grandiflorum* and *H. umbratica*. Collectively, the three species share 1,816 gene families. The shared gene families among the three species exhibit only two significant GO enrichment: one related to pollen recognition (GO:0048544) and the other associated with protein localization to the cell surface (GO:0034394). Among the exclusive and shared gene families and singletons, many are linked to fruit quality, maturation, development of organoleptic characteristics, general plant development, and resistance to pathogens (Figure 5 B and Tables S7 and S8).

**Figure 5:**
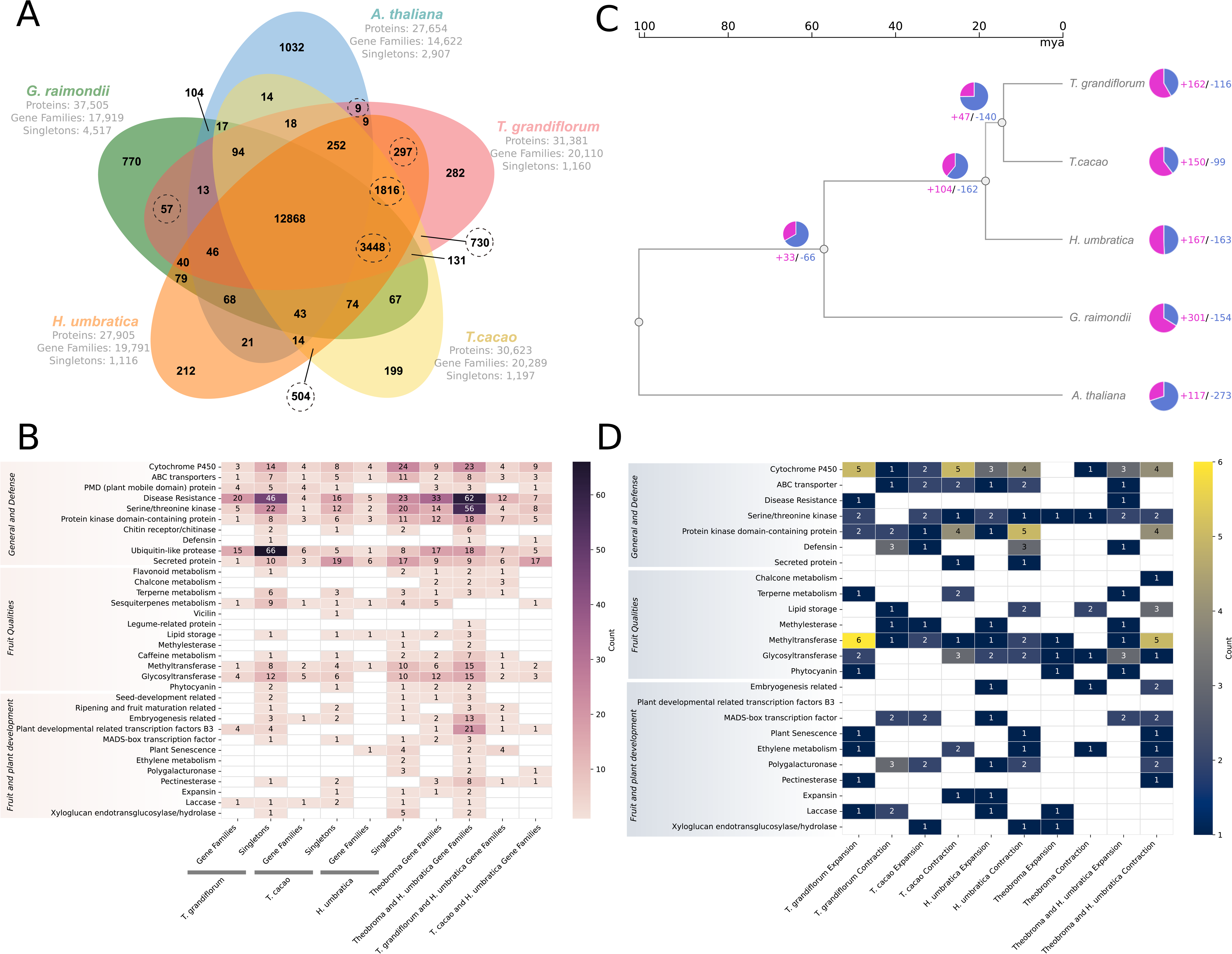
Comparative Analyses Across Malvaceae Species Focusing on Functions Related to Plant Differentiation, Fruit and Seed Development, and Organoleptic and Physicochemical Qualities. **A.** A Venn diagram illustrates the shared and exclusive orthologous clusters (gene families) identified across four Malvaceae species and *Arabidopsis thaliana*. **B.** The identification of gene families and singletons encompasses a range of functions with predicted roles in various aspects of plant and fruit development. These include Cytochrome P450 and ABC transporters, which are pivotal in synthesizing secondary metabolites and nutrient uptake, respectively, influencing plant growth and fruit quality. Plant Mobile Domain (PMD) proteins and disease resistance genes play roles in stress response and plant health, indirectly impacting fruit quality. Serine/threonine kinase, protein kinase domain-containing proteins, and several metabolism-related genes (flavonoid, chalcone, terpene, sesquiterpenes) regulate pathways critical for plant growth, development, and the organoleptic properties of fruits. Genes related to defense mechanisms (chitin receptor/chitinase, defensin, ubiquitin-like protease) and cell wall composition (methylesterase, polygalacturonase, pectinesterase, expansin, laccase, xyloglucan endotransglucosylase/hydrolase) are also identified, reflecting their roles in maintaining plant health and influencing fruit texture and firmness. Furthermore, genes involved in seed development (vicilin, legume-related protein, lipid storage) and various transcription factors (including MADS-box) are noted for their influence on plant growth and developmental processes. **C.** A phylogenetic tree delineates the evolutionary timeline of the Malvaceae species with *A. thaliana* serving as the outgroup. An accompanying pie chart displays the proportions of gene families that have expanded or contracted, indicating evolutionary dynamics. **D.** The analysis of expanded and contracted gene families focuses on their common functions and roles, as detailed in section **B**, shedding light on the evolutionary adaptations of these species.

Moreover, the analysis of gene expansion and contraction revealed distinct patterns across the analyzed Malvaceae (Figure 5 C). Despite the GO enrichment analyses did not indicate any other statistically significant enrichment, we were able to determine specific gene functions related to important agronomical traits indicating groups of gene families that were expanded and contracted in each species (Figure 5 D and Table S9).

We found that unique profiles of singletons and gene families (both expanded and contracted) are primarily present in cytochrome P450, ABC transporters, and other functions related to plant development and pathogen defense. This indicates specific adaptations and responses to domestication, environmental changes, and various stresses. For instance, numerous gene families and singletons genes belonging the PMD domain-containing protein identified uniquely in *T. grandiflorum* and *T. cacao* likely plays a role in developmental control [69], while the singletons genes encoding to chitin receptor/chitinase (i.e. TgrandC1074G00000003550 and TgrandC1074G00000000752) may be crucial for fungal resistance.

Exclusive gene profiles linked to fruit and seed quality, notably in lipid storage and secondary metabolite functions were identified (Figure 5 B and D). The storage lipids in seeds are key components of the quality of cocoa butter and chocolate in cacao and cupulate and cosmetics products in cupuassu [11,70]. Additionally, unique gene profiles involved in flavonoid, chalcone, terpenoid, and sesquiterpene metabolism might contribute to the distinct aromas of cacao and cupuassu seeds. Furthermore, different profiles in purine alkaloid metabolism could explain the flavor differences between both *Theobroma* species

Moreover, distinct patterns of enzymes such as methyltransferase, glycosyltransferase, and phytocyanin were identified, all crucial to secondary metabolism and linked to fruit traits. Methyltransferases are key in secondary metabolite metabolism (phenylpropanoids, flavonoids, alkaloids) affecting flavor, pulp, and peel color [71–73]. Glycosyltransferases, catalyzing glycosylation reactions for various substrates, including plant hormones and secondary metabolites, influence fruit ripening and seed development [74,75]. Additionally, the unique gene pattern of phytocyanin, involved in growth and stress resilience [76], may be linked to the adaptability in challenging environmental conditions.

### Spatial Arrangement and Distribution of Duplicated Genes Reveals Evolutionary Insights into Fruit and Seed Quality and Defense Mechanism Origins

The spatial gene arrangement and distribution in the genome of the three species was evaluated by comprehensive GO enrichment analyses (Figure 6 and Table S10). This analysis revealed distinct functional variations across various types of gene duplications, including WGD events, and tandem, proximal, dispersed, and singleton duplicates.

**Figure 6:**
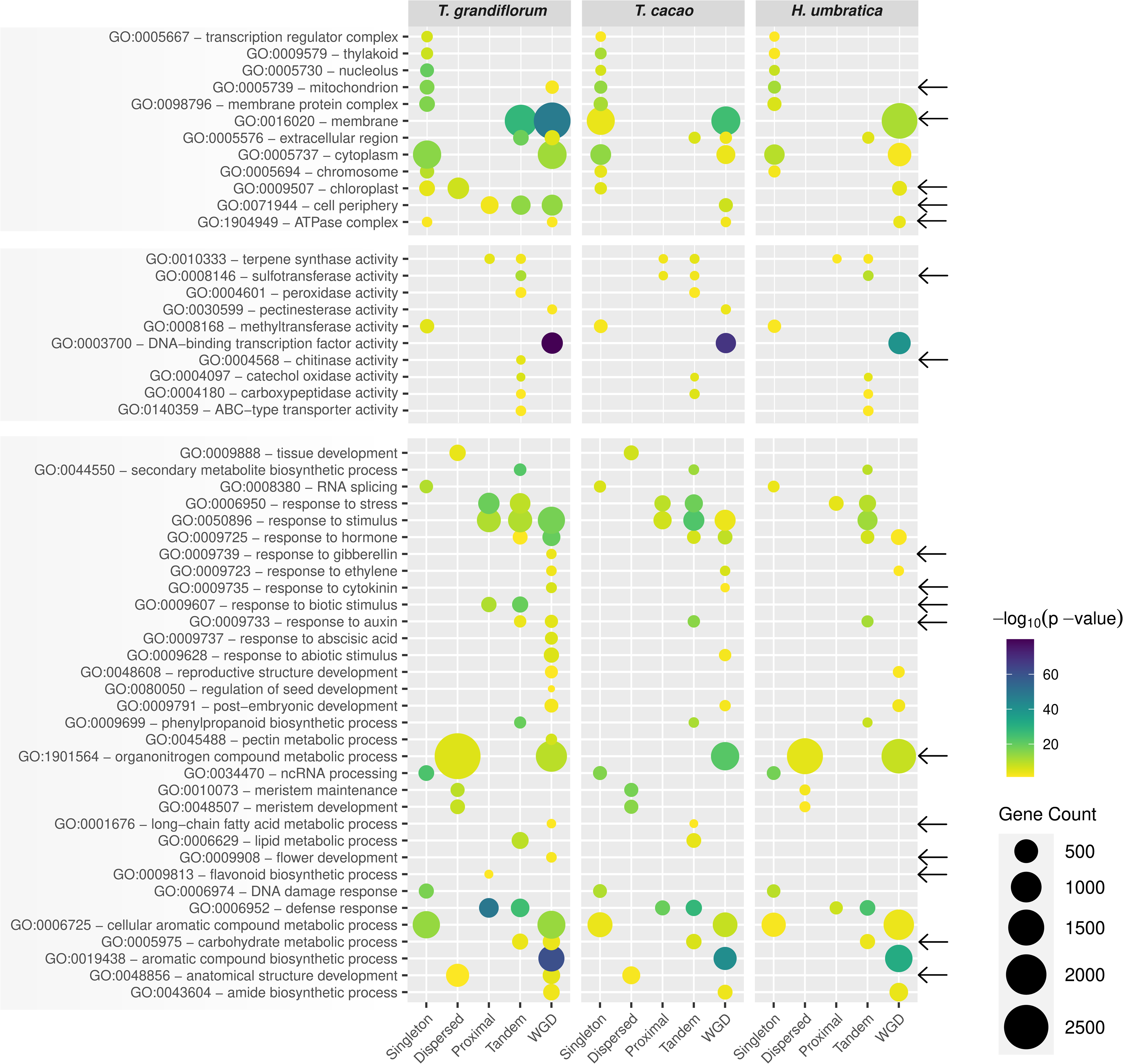
Gene Ontology Enrichment and Comparative Analysis Across *Theobroma grandiflorum*, *T. cacao*, and *Herrania umbratica*. Black arrows highlight GO terms that are exclusively enriched in *T. grandiflorum*, either in duplicated genes or singletons. These terms provide insights into the unique biological processes, cellular components, and molecular functions connected with fruit and seed quality and defense mechanism that are particularly prominent in *T. grandiflorum* compared to the other species.

#### Cellular Component Ontology Variations

1. *T. grandiflorum* uniquely exhibited enrichment in WGD-derived genes associated with the mitochondrion (GO:0005739) and tandem genes linked to the membrane (GO:0016020). In contrast, *T. cacao* showed an enrichment of membrane-associated (GO:0016020) singleton genes. Additionally, *T. grandiflorum* also showed enrichment in chloroplast (GO:0009507) in dispersed duplicates, and cell periphery (GO:0071944) in proximal and tandem duplicated genes, with an exclusive enrichment of ATPase complex (GO:1904949) in singleton genes.

#### Molecular Function Ontology Variations

Notably, sulfotransferase activity (GO:0008146), potentially influencing flavonoid metabolism [77], was enriched in both proximal and tandem duplicates of *T. caca*o. In contrast, this activity was enriched exclusively in tandem duplicates in the cupuassu genome. Chitinase activity (GO:0004568), likely linked to defense against fungal pathogens [21], was enriched only in *T. grandiflorum* tandem duplicated genes.

#### GO Terms Related to Fruit and Seed Traits

Numerous GO terms potentially related to fruit and seed traits were identified as enriched in duplicated genes. These included terpene synthase activity (GO:0010333) and the flavonoid biosynthetic process (GO:0009813), which were enriched in proximal duplicated genes of *T. grandiflorum*. Additionally, GO terms associated with secondary metabolite biosynthesis, lipid metabolism, phenylpropanoid biosynthesis, catechol oxidase activity, and carboxypeptidase activity were predominantly enriched in tandem genes. Notably, GO terms related to the organonitrogen compound metabolic process (GO:1901564) and long-chain fatty acid metabolic process (GO:0001676) showed diverse enrichment patterns across species. This finding is particularly noteworthy due to the distinct differences in fatty acid composition between cacao seeds and cupuassu fruit pulp. Specifically, cacao seeds exhibit a higher concentration of saturated fatty acids, predominantly palmitic and stearic acids, followed by desaturated fatty acids, including oleic and linoleic acids [78]. In contrast, cupuassu is characterized by a richness in desaturated fatty acids [79].

#### GO Terms Related to Fruit Aroma and Ripening Process

The cellular aromatic compound metabolic process (GO:0006725), which may affect fruit aroma and plant defense [80], was enriched in singletons and WGD-derived genes in all three species. Pectinesterase activity (GO:0030599), potentially related to fruit ripening and cell wall fortification [81], was enriched in WGD-derived genes of *T. grandiflorum* and *T. cacao*, but not in *H. umbratica*.

#### GO Terms Related to Fruit Morphology and Hormonal Responses

Genes associated with meristem development (GO:0048507/GO:0010073) and anatomical structure development (GO:0048856) were predominantly enriched in dispersed duplicates. In contrast, genes related to seed development (GO:0080050) and flower development (GO:0009908) showed enrichment in WGD-derived genes. These WGD-derived genes were also enriched in terms related to hormonal responses and DNA binding transcription factor activity (GO:0003700), while singleton genes showed enrichment in ncRNA processing (GO:0034470), RNA splicing (GO:0008380), and DNA damage responses (GO:0006974).

#### GO Terms Related to Defense Responses and Stress Reactions

Genes involved in the defense response (GO:0006952) and response to stress (GO:0006950) were enriched in proximal and tandem repeated genes. The response to biotic stimulus (GO:0009607) was also enriched in these gene types, whereas the response to abiotic stimulus (GO:0009628) was more prevalent in WGD-derived genes.

### Positively Selected Retained Dispersed, Proximal and Tandem Duplications: Potential Drivers of Fruit and Pathogen Resistance Evolution ?

From a general evolutionary perspective, genes derived from WGD events are typically ancient and often well-integrated into the existing genetic framework, which allows ample time to functionally diverge [82]. In contrast, genes from tandem, proximal, and dispersed duplications are generally younger, often emerging in response to environmental challenges and stressors [83,84], and possibly influenced by domestication process. In parallel, singleton genes, often originating from genome fractionation events after WGD, play crucial roles in core cellular functions and essential physiological processes [85–87].

During evolutionary timeframes and through domestication processes, new genes were created by duplication and were lost over time. Interestingly, some duplicated genes are retained and can acquire new roles (neofunctionalization) or specialize in aspects of their original function (subfunctionalization), contributing to morphological innovations and the development of new functionalities, including the enhancement of disease resistance, and increased stress adaptability [86,88].

To contextualize these evolutionary processes, we evaluated the rate of non-synonymous nucleotide substitutions (Ka) relative to synonymous substitutions (Ks) across different gene duplication types (Figure 7A). A significant majority of duplicated genes in *H. umbratica* (97.39%), *T. cacao* (95.37%), and *T. grandiflorum* (93.85%) are under purifying selection, a trend consistent with observations in other plant species [43]. WGD-derived genes in all species exhibit strong purifying selection with a mean Ka/Ks of 0.132. Dispersed duplicates largely follow this trend (mean Ka/Ks of 0.165), with occasional peaks suggesting a balance between purifying and positive selection.

**Figure 7:**
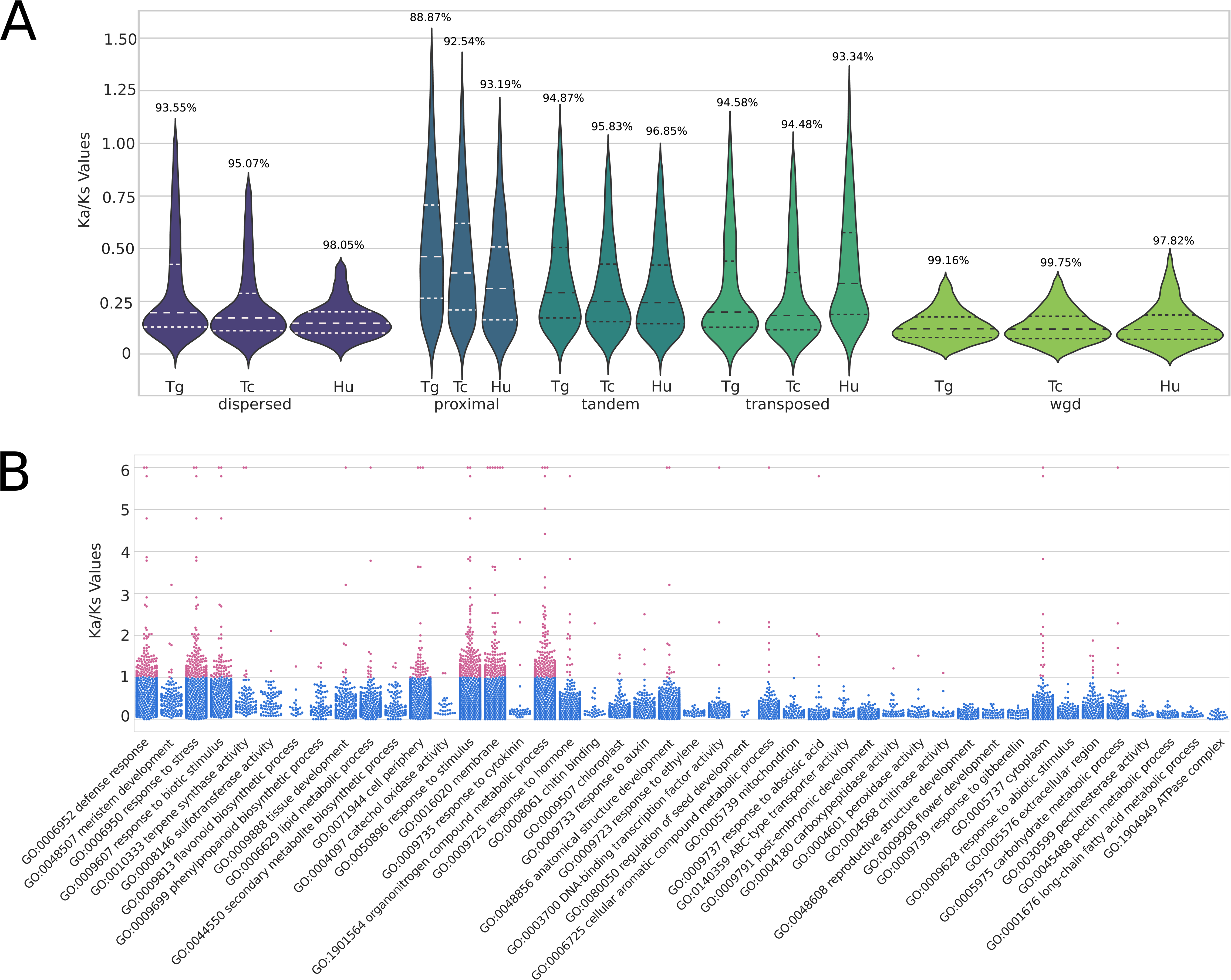
Positive Selection Analysis of Duplicated Genes in *Theobroma grandiflorum*, *T. cacao*, and *Herrania umbratica.* **A.** A violin plot displays the distribution of the Ka/Ks ratios for gene pairs resulting from dispersed, proximal, and tandem duplication in *Theobroma grandiflorum*, *T. cacao*, and *Herrania umbratica*. The number above each plot indicates the percentage of duplicated genes under purifying selection. B. A swarmplot illustrates the Ka/Ks ratio distributions for selected Gene Ontology (GO) terms associated with fruit traits and defense mechanisms in *T. grandiflorum*. This plot provides insights into the selective pressures acting on genes related to these specific functions.

Although the majority of proximal (mean Ka/Ks = 0.444), tandem (mean Ka/Ks = 0.331), and transposed (mean Ka/Ks = 0.329) gene pairs demonstrate a trend to be under purifying selection, there is an evident trend towards greater tolerance to variation. These findings support the hypothesis of post-speciation adaptation in these gene groups, likely linked to diversification or domestication influences.

Indeed, upon detailed examination, a significant portion of duplicated genes in *T. grandiflorum* (6.15%), *T. cacao* (4.62%), and *H. umbratica* (2.6%)—associated with GO terms related to plant defense, fruit and seed traits—were found to be under positive selection (Figure 7 B, Figure S6, Table S11).

For instance, in the evolutionary battle between plants and their adversaries, defense-related genes often undergo positive selection [89,90]. This is exemplified by several clusters of tandemly duplicated genes linked to defense responses and plant disease resistance, which demonstrate strong positive selection in *T. grandiflorum* (139 genes), *T. cacao* (40 genes), and *H. umbratica* (17 genes). Notably, in *T. grandiflorum*, a significant concentration of these genes is found in chromosomes 6, 7, and 10 (Figure S7).

A cluster of genes on chromosome 6 of *T. grandiflorum* corresponds with an identified cupuassu WBD-resistance quantitative trait locus (QTL) [18]. Within this QTL, the *TgPR3* gene, encoding a chitinase, was associated with WBD resistance [21]. The cupuassu genome we sequenced displays the chitinase gene (TgrandC1074G00000024418), which is encircled by a multitude of disease resistance genes located within this QTL. Some of these disease resistance genes are tandemly duplicated and exhibit signs of positive selection, suggesting a robust assembly of disease resistance genes in this specific QTL (Table S12). However, it is essential to recognize that the cupuassu genome under analysis is from a *Moniliophthora perniciosa*-susceptible genotype. As a result, the evolutionary gene pattern identified may not necessarily confer resistance to WBD, but could potentially be associated with resistance to other pathogens.

In the terpene synthase activity (GO:0010333), tandem arrays encoding a number of delta-cadinene synthase are under positive selection across the three species. This enzyme plays a role in sesquiterpene biosynthesis, crucial for plant defense and the production of compounds like gossypol in cotton seeds [91,92]. It was also considered a key candidate for studying cacao-insect resistance interplay [11]. Interestingly, *T. grandiflorum* uniquely harbors tandem repeated genes encoding a probable terpene synthase (TgrandC1074G00000007568 and TgrandC1074G00000007569), hinting at regulatory role in terpenoid biosynthesis with potential ramifications for fruit aroma and flavor. Conversely, *T. cacao* possesses positively selected tandem repeated genes encoding a potential nerolidol synthase (Tcacao-CriolloG00000024422 and Tcacao-CriolloG00000024423). In cacao, this enzyme contributes to linalool biosynthesis, producing volatile compounds that belong to the monoterpenes group. These compounds are abundant in cocoa beans and are responsible for their floral aroma [93]. In grapes, this enzyme enhances the aroma of certain varieties [94]. Additionally, in rice, it is associated with the production of an antibacterial compound effective against bacterial pathogens [95].

Within the flavonoid biosynthetic pathway (GO:0009813) of *T. grandiflorum*, a gene encoding a positively selected tandem duplicated naringenin 2-oxoglutarate 3-dioxygenase (TgrandC1074G00000004751 and TgrandC1074G00000004753) may emerges as pivotal in specific flavonoid, anthocyanidins, catechins and proanthocyanidins biosynthesis. Given naringenin documented broad-spectrum biological impacts on human health [96], it is conceivable that this gene plays a role in the distinct antioxidant properties of cupuassu [97], further influencing the fruit unique taste and aroma.

Another noteworthy instance of tandemly duplicated genes under positive selection, potentially linked to fruit and seed characteristics, involves those engaged in the lipid metabolic process (GO:0006629). Both cupuassu and cacao present distinct pattern, possibly related to their unique seed properties. Specifically, cupuassu has a positively selected and tandemly duplicated gene related to lipid storage in fruits, known as patatin (TgrandC1074G00000017909 and TgrandC1074G00000017911). Originally identified in potato (*Solanum tuberosum L.*) tubers, patatin is renowned for its antioxidant potential [98] and its exceptional nutritional value, making it an appealing food additive due to its solubility and emulsifying properties [99,100].

In contrast, *T. cacao* features a tandem duplicated phospholipase A1 positively selected (Tcacao-CriolloG00000024071 and Tcacao-CriolloG00000024072), which could modulate the fruit phospholipid profile. For instance, this phospholipase may be involved in linoleic acid metabolism [99–101], central to the production of desaturated fatty acids present in cacao-derived chocolates [78]. Meanwhile, *T. grandiflorum*, displays a tandem repeated gene encoding a fatty acyl-CoA reductase enzyme (TgrandC1074G00000003252 and TgrandC1074G00000003253), potentially influencing the lipid content and composition of seeds, impacting wax biosynthesis [102] and, by extension, the fruit cuticle, water retention, and shelf life.

Furthermore, *T. cacao* possesses two dispersed duplicated and positively selected pectinesterases (Tcacao-CriolloG00000016685 and Tcacao-CriolloG00000021823) that might play a significant role in the ripening of cacao fruit. Interestingly, neither *T. grandiflorum* nor *H. umbratica* exhibit positively selected pectinesterases. This observation may be associated with the behavior of cacao tree fruits, which do not fall when ripe but remain attached to the tree until manually harvested. [67].

## Conclusions

Recent advancements in long-read sequencing, chromatin interaction technologies, and comparative genomics have significantly enriched our understanding of genome evolution, particularly in the *Theobroma* genus, and have contributed to insights into phenotypic variation [13,103]. These tools facilitate in-depth analysis of plant development and the determinants of disease resistance, offering substantial biotechnological implications. They are becoming increasingly essential in crop breeding to address challenges such as climate change and food security.

Our study presents a chromosome-scale genome assembly of *T. grandiflorum*, enhancing genetic resources for breeding and sustainable horticulture. We have uncovered evolutionary insights into the origins of genes linked to key agronomic traits. Furthermore, we identified unique gene families and singletons in Malvaceae species, which may be instrumental in organs development, defense, adaptation, and distinctive fruit traits. The variations in gene presence and absence (and gene family expansion and contraction) among these species might be linked to unique mechanisms of gene retention and loss, which in turn are closely related to the generation of phenotypic diversity and innovation [104]. Concurrently, we revealed that many retained duplicated genes related to plant defense, fruit, and seed production are under positive selection. This finding also aligns with known processes of phenotypic novelty emergence, leading to speciation and diversification [86,105,106]. By providing a comprehensive candidate genes list, we aim not only to support breeding initiatives but also to deepen our understanding of the cupuassu genome biology. We believe that the results presented here lay the groundwork for advanced functional genomic interventions and tailored cultivation methods. This could potentially enhance species conservation and farmer productivity, thereby further impacting the Amazonian bioeconomy. In conclusion, our findings offer valuable insights into the unique evolutionary pathways and domestication of *T. grandiflorum* and *T. cacao*, particularly in terms of pathogen resistance, fruit and seed development and adaptive strategies post-diversification.

## Abbreviations

BUSCO: Benchmarking Universal Single-Copy Orthologs
CDS: coding sequence
CTAB: Cetyltrimethylammonium Bromide
FP: Frosty pod
GO: Gene Ontology
HiC: Chromosome conformation capture techniques
HMW: High molecular weight
Ka: non-synonymous nucleotide substitutions
Ks: synonymous substitutions
LAI: LTR Assembly Index
LARD: Large Retrotransposon Derivatives
lncRNAs: long non-coding RNAs
LTR-RT: Long Terminal Repeat Retrotransposons
miRNAs: microRNAs
mya: million years ago
QTL: Quantitative trait locus
rDNA: ribosomal DNA
snoRNAs: small nucleolar RNAs
snRNAs: small nuclear RNAs
sRNAs: small RNAs
T2T: telomere-to-telomere
TE: Transposable Elements
TRIM: Terminal-repeat Retrotransposons in Miniature
tRNAs: transfer RNAs
WBD: Witches’ broom disease
WGD: Whole-genome duplication

## Author Contributions

**Rafael Moysés Alves**: Conceptualization; Data curation; project administration; writing—original draft; writing—review and editing. **Vinicius A. C. de Abreu**: Conceptualization; Data curation; formal analysis; software; investigation; supervision; project administration; writing—review and editing. **Rafaely Pantoja Oliveira**: Formal analysis; investigation; review and editing. **João Victor dos Anjos Almeida**: Formal analysis; investigation; review and editing. **Mauro de Medeiros de Oliveira**: Resources; review and editing. **Saura R. Silva**: Resources; review and editing. **Alexandre R. Paschoal:** Investigation; methodology; software, review and editing. **Sintia Almeida:** investigation; review and editing. **Pedro A. F. de Souza:** investigation; review and editing. **Jesus A. Ferro**: resources; review and editing. **Vitor F. O. Miranda**: Resources, review and editing. **Antonio Figueira**: Investigation; writing—review and editing. **Douglas S. Domingue**s: Investigation; writing—review and editing. **Alessandro M. Varani**: Conceptualization; Data curation; formal analysis; software; investigation; methodology; resources; project administration; supervision; writing—review and editing.

## Acknowledgments

We thank the Arizona Genomics Institute (Tucson, AZ – USA) for providing all the HMW DNA extraction and PacBio Sequel IIe sequencing support. This study was financed by Fundação de Amparo à Pesquisa do Estado do São Paulo – FAPESP, grant #2019/25176-0 to AMV, and Fundação Amazônia de Amparo a Estudos e Pesquisas – FAPESPA, grant #075/2020 to VACA and RMA. Fundação Araucária supported ARP in NAPI Bioinformática project grant #66.2021. These funding agencies had no role in study design, the collection, analysis, and interpretation of data, or manuscript writing.

## SUPPLEMENTAL MATERIAL

**Supplementary Information 1.** HMW DNA extraction, Sequencing QC, Bioinformatics procedures used to annotate *Theobroma grandiflorum, T. cacao* and *Herrania umbratica* genomes, and additional notes.

**Figure S1. Distribution of the ncRNA identified in *Theobroma grandiflorum* genome.**

**Figure S2.** LTR insertion time of Gypsy and Copia elements. **A.** *Theobroma grandiflorum*, **B.** *T. cacao*, and **C.** *Herrania umbratica*. The vertical black line represents the median, and the dotted line represents the mean.

**Figure S3.** TE_density analyses of all *Theobroma grandiflorum* chromosomes.

**Figure S4. A.** Microsynteny and coliniarity example of subtelomeric regions of *T. grandiflorum*, *T. cacao* and *H. umbratica*, **B.** Microsynteny and coliniarity example of pericentromeric regions of *T. grandiflorum*, *T. cacao* and *H. umbratica*. Blue represents genes in the forward direction, green indicates genes in the reverse direction, and orange denotes transposable elements (TEs).

**Figure S5.** Alignment of GEX1 gene from CH4 loci generated on Jalview (Procter et al., 2021).

**Figure S6.** Box-plot and swarmplot showing the the Ka/Ks ratio distributions of the selected GO terms associated with fruit traits and defense mechanisms. A. *Theobroma cacao*, B. *Herrania umbratica*.

**Figure S7.** Genomic mapping of plant disease resistant genes in *Theobroma grandiflorum* chromosomes. Genes under positive selection are shown in red. The cupuassu WBD-resistant QTL is shown in blue.

**Table S1.** GenBank SRA accession numbers used for transcriptome assembly. **A.** All *Theobroma cacao* RNAseq data used. **B.** *Herrania umbratica* RNAseq data used.

**Table S2.** Genome assembly statistics and completeness scores of the three Malvaceae.

**Table S3.** Genome annotation features and statistics of the three Malvaceae.

**Table S4.** Retrocopies identified in *Theobroma grandiflorum*, *T. cacao*, and *Herrania umbratica*, with associated raw data.

**Table S5.** Genome structural features and Statistics for each *Theobroma grandiflorum* chromosome.

**Table S6.** Tranposable elements summary table and statistics identified in *Theobroma grandiflorum*, *T. cacao*, and *Herrania umbratica*.

**Table S7.** Exclusive gene families identified for each genome analyzed.

**Table S8.** Singletons identified in each genome analyzed.

**Table S9.** Expanded and Contracted gene families identified in each genome analyzed.

**Table S10.** GO enrichment analyses raw data.

**Table S11.** Genes and GO terms identified as positively selected by Ka/Ks analysis.

**Table S12.** Gene content and features of cupuassu WBD-resistance QTL.

## REFERENCES

1. Cuatrecasas J. Cacao and Its Allies: A Taxonomic Revision of the Genus Theobroma. Smithsonian Inst;

2. The Angiosperm Phylogeny Group. An update of the Angiosperm Phylogeny Group classification for the orders and families of flowering plants: APG IV. Bot J Linn Soc. 2016; doi: 10.1111/boj.12385.

3. da Silva RA, Souza G, Lemos LSL, Lopes UV, Patrocínio NGRB, Alves RM, et al.. Genome size, cytogenetic data and transferability of EST-SSRs markers in wild and cultivated species of the genus Theobroma L. (Byttnerioideae, Malvaceae). PLoS One. 2017; doi: 10.1371/journal.pone.0170799.

4. Freitas ÍR, Pirani JR, Colli-Silva M. CACAU PARA QUÊ? LEVANTAMENTO BIBLIOGRÁFICO SOBRE OS USOS MATERIAIS E SIMBÓLICOS DAS ESPÉCIES DE CACAUS DO BRASIL. Ethnoscientia - Brazilian Journal of Ethnobiology and Ethnoecology. 2023; doi: 10.18542/ethnoscientia.v8i1.12940.

5. Garcia TB, Potiguara RC de V, Kikuchi TYS, Demarco D, Aguiar-Dias ACA de. Leaf anatomical features of three Theobroma species (Malvaceae s.l.) native to the Brazilian Amazon. Acta Amaz. Instituto Nacional de Pesquisas da Amazônia; 2014; doi: 10.1590/1809-4392201300653.

6. Colli-Silva M, Richardson JE, Neves EG, Watling J, Figueira A, Pirani JR. Domestication of the Amazonian fruit tree cupuaçu may have stretched over the past 8000 years. Commun Earth Environ. Nature Publishing Group; 2023; doi: 10.1038/s43247-023-01066-z.

7. Alves RM, Chaves SF da S. Selection of Theobroma grandiflorum clones adapted to agroforestry systems using an additive index. Acta Scientiarum Agronomy. 2023; doi: 10.4025/actasciagron.v45i1.57519.

8. Alves RM, Chaves SF da S. BRS Careca, BRS Fartura, BRS Duquesa, BRS Curinga, and BRS Golias: new cupuassu tree cultivars. Crop Breed Appl Biotechnol. Crop Breeding and Applied Biotechnology; 2020; doi: 10.1590/1984-70332020v20n4c66.

9. Leal GA, Albuquerque PSB, Figueira A. Genes differentially expressed in Theobroma cacao associated with resistance to witches’ broom disease caused by Crinipellis perniciosa. Mol Plant Pathol. 2007; doi: 10.1111/j.1364-3703.2007.00393.x.

10. Falcão LL, Silva-Werneck JO, Albuquerque PSB, Alves RM, Grynberg P, Togawa RC, et al.. Comparative transcriptomics of cupuassu (Theobroma grandiflorum) offers insights into the early defense mechanism to Moniliophthora perniciosa, the causal agent of witches’ broom disease. Journal of Plant Interactions. Taylor & Francis; 2022; doi: 10.1080/17429145.2022.2144650.

11. Argout X, Salse J, Aury J-M, Guiltinan MJ, Droc G, Gouzy J, et al.. The genome of Theobroma cacao. Nat Genet. 2011; doi: 10.1038/ng.736.

12. Argout X, Martin G, Droc G, Fouet O, Labadie K, Rivals E, et al.. The cacao Criollo genome v2.0: an improved version of the genome for genetic and functional genomic studies. BMC Genomics. 2017; doi: 10.1186/s12864-017-4120-9.

13. Argout X, Droc G, Fouet O, Rouard M, Labadie K, Rhoné B, et al.. Pangenomic exploration of Theobroma cacao: New Insights into Gene Content Diversity and Selection During Domestication. bioRxiv. Cold Spring Harbor Laboratory; 2023; doi: 10.1101/2023.11.03.565324.

14. Motamayor JC, Mockaitis K, Schmutz J, Haiminen N, Livingstone D, Cornejo O, et al.. The genome sequence of the most widely cultivated cacao type and its use to identify candidate genes regulating pod color. Genome Biol. 2013; doi: 10.1186/gb-2013-14-6-r53.

15. Morrissey J, Stack JC, Valls R, Motamayor JC. Low-cost assembly of a cacao crop genome is able to resolve complex heterozygous bubbles. Hortic Res. Nature Publishing Group; 2019; doi: 10.1038/s41438-019-0125-7.

16. Hämälä T, Wafula EK, Guiltinan MJ, Ralph PE, dePamphilis CW, Tiffin P. Genomic structural variants constrain and facilitate adaptation in natural populations of Theobroma cacao, the chocolate tree. Proc Natl Acad Sci U S A. 2021; doi: 10.1073/pnas.2102914118.

17. Colli-Silva M, Richardson J, Pirani J. A taxonomic dataset of preserved specimen occurrences of Theobroma and Herrania (Malvaceae, Byttnerioideae) stored in 2020. Biodiversity Data Journal. Pensoft Publishers; 2023; doi: 10.3897/BDJ.11.e99646.

18. Mournet P, de Albuquerque PSB, Alves RM, Silva-Werneck JO, Rivallan R, Marcellino LH, et al.. A reference high-density genetic map of Theobroma grandiflorum (Willd. ex Spreng) and QTL detection for resistance to witches’ broom disease (Moniliophthora perniciosa). Tree Genetics & Genomes. 2020; doi: 10.1007/s11295-020-01479-3.

19. Niu Y-F, Ni S-B, Liu J. The complete chloroplast genome of Theobroma grandiflorum, an important tropical crop. Mitochondrial DNA B Resour. 2019; doi: 10.1080/23802359.2019.1693291.

20. de Abreu VAC, Moysés Alves R, Silva SR, Ferro JA, Domingues DS, Miranda VFO, et al.. Comparative analyses of Theobroma cacao and T. grandiflorum mitogenomes reveal conserved gene content embedded within complex and plastic structures. Gene. 2023; doi: 10.1016/j.gene.2022.146904.

21. Santana Silva RJ, Alves RM, Peres Gramacho K, Marcellino LH, Micheli F. Involvement of structurally distinct cupuassu chitinases and osmotin in plant resistance to the fungus Moniliophthora perniciosa. Plant Physiol Biochem. 2020; doi: 10.1016/j.plaphy.2020.01.009.

22. Doyle JJ, Doyle JL, editors. A rapid DNA isolation procedure for small quantities of fresh leaf tissue. PHYTOCHEMICAL BULLETIN.

23. Ranallo-Benavidez TR, Jaron KS, Schatz MC. GenomeScope 2.0 and Smudgeplot for reference-free profiling of polyploid genomes. Nat Commun. 2020; doi: 10.1038/s41467-020-14998-3.

24. Kokot M, Dlugosz M, Deorowicz S. KMC 3: counting and manipulating k-mer statistics. Bioinformatics. 2017; doi: 10.1093/bioinformatics/btx304.

25. Cheng H, Concepcion GT, Feng X, Zhang H, Li H. Haplotype-resolved de novo assembly using phased assembly graphs with hifiasm. Nat Methods. 2021; doi: 10.1038/s41592-020-01056-5.

26. Wood DE, Lu J, Langmead B. Improved metagenomic analysis with Kraken 2. Genome Biology. 2019; doi: 10.1186/s13059-019-1891-0.

27. Li H, Durbin R. Fast and accurate short read alignment with Burrows–Wheeler transform. Bioinformatics. 2009; doi: 10.1093/bioinformatics/btp324.

28. Durand NC, Shamim MS, Machol I, Rao SSP, Huntley MH, Lander ES, et al.. Juicer Provides a One-Click System for Analyzing Loop-Resolution Hi-C Experiments. Cell Syst. 2016; doi: 10.1016/j.cels.2016.07.002.

29. Dudchenko O, Batra SS, Omer AD, Nyquist SK, Hoeger M, Durand NC, et al.. De novo assembly of the Aedes aegypti genome using Hi-C yields chromosome-length scaffolds. Science. American Association for the Advancement of Science; 2017; doi: 10.1126/science.aal3327.

30. Zimin AV, Puiu D, Luo M-C, Zhu T, Koren S, Marçais G, et al.. Hybrid assembly of the large and highly repetitive genome of Aegilops tauschii, a progenitor of bread wheat, with the MaSuRCA mega-reads algorithm. Genome Res. 2017; doi: 10.1101/gr.213405.116.

31. Zhou C, McCarthy SA, Durbin R. YaHS: yet another Hi-C scaffolding tool. Bioinformatics. 2023; doi: 10.1093/bioinformatics/btac808.

32. Rhie A, Walenz BP, Koren S, Phillippy AM. Merqury: reference-free quality, completeness, and phasing assessment for genome assemblies. Genome Biol. 2020; doi: 10.1186/s13059-020-02134-9.

33. Ou S, Chen J, Jiang N. Assessing genome assembly quality using the LTR Assembly Index (LAI). Nucleic Acids Res. 2018; doi: 10.1093/nar/gky730.

34. Kriventseva EV, Kuznetsov D, Tegenfeldt F, Manni M, Dias R, Simão FA, et al.. OrthoDB v10: sampling the diversity of animal, plant, fungal, protist, bacterial and viral genomes for evolutionary and functional annotations of orthologs. Nucleic Acids Res. 2019; doi: 10.1093/nar/gky1053.

35. Manni M, Berkeley MR, Seppey M, Zdobnov EM. BUSCO: Assessing Genomic Data Quality and Beyond. Curr Protoc. 2021; doi: 10.1002/cpz1.323.

36. Haas BJ, Papanicolaou A, Yassour M, Grabherr M, Blood PD, Bowden J, et al.. De novo transcript sequence reconstruction from RNA-seq using the Trinity platform for reference generation and analysis. Nat Protoc. Nature Publishing Group; 2013; doi: 10.1038/nprot.2013.084.

37. Kim D, Paggi JM, Park C, Bennett C, Salzberg SL. Graph-based genome alignment and genotyping with HISAT2 and HISAT-genotype. Nat Biotechnol. Nature Publishing Group; 2019; doi: 10.1038/s41587-019-0201-4.

38. Li H. Minimap2: pairwise alignment for nucleotide sequences. Bioinformatics. 2018; doi: 10.1093/bioinformatics/bty191.

39. Kovaka S, Zimin AV, Pertea GM, Razaghi R, Salzberg SL, Pertea M. Transcriptome assembly from long-read RNA-seq alignments with StringTie2. Genome Biology. 2019; doi: 10.1186/s13059-019-1910-1.

40. Haas BJ, Delcher AL, Mount SM, Wortman JR, Smith RK, Hannick LI, et al.. Improving the Arabidopsis genome annotation using maximal transcript alignment assemblies. Nucleic Acids Res. 2003; doi: 10.1093/nar/gkg770.

41. Vuruputoor VS, Monyak D, Fetter KC, Webster C, Bhattarai A, Shrestha B, et al.. Welcome to the big leaves: Best practices for improving genome annotation in non-model plant genomes. Appl Plant Sci. 2023; doi: 10.1002/aps3.11533.

42. Wang Y, Jia L, Tian G, Dong Y, Zhang X, Zhou Z, et al.. shinyCircos-V2.0: Leveraging the creation of Circos plot with enhanced usability and advanced features. iMeta. 2023; doi: 10.1002/imt2.109.

43. Qiao X, Li Q, Yin H, Qi K, Li L, Wang R, et al.. Gene duplication and evolution in recurring polyploidization–diploidization cycles in plants. Genome Biology. 2019; doi: 10.1186/s13059-019-1650-2.

44. Wang Y, Tang H, Debarry JD, Tan X, Li J, Wang X, et al.. MCScanX: a toolkit for detection and evolutionary analysis of gene synteny and collinearity. Nucleic Acids Res. 2012; doi: 10.1093/nar/gkr1293.

45. Bandi V, Gutwin C, Siri JN, Neufeld E, Sharpe A, Parkin I. Visualization Tools for Genomic Conservation. Methods Mol Biol. 2022; doi: 10.1007/978-1-0716-2067-0_16.

46. Tang H, Bowers JE, Wang X, Ming R, Alam M, Paterson AH. Synteny and Collinearity in Plant Genomes. Science. 2008; doi: 10.1126/science.1153917.

47. Chao J, Li Z, Sun Y, Aluko OO, Wu X, Wang Q, et al.. MG2C: a user-friendly online tool for drawing genetic maps. Mol Hortic. 2021; doi: 10.1186/s43897-021-00020-x.

48. Teresi SJ, Teresi MB, Edger PP. TE Density: a tool to investigate the biology of transposable elements. Mobile DNA. 2022; doi: 10.1186/s13100-022-00264-4.

49. Mauri M, Elli T, Caviglia G, Uboldi G, Azzi M. RAWGraphs: A Visualisation Platform to Create Open Outputs. Proceedings of the 12th Biannual Conference on Italian SIGCHI Chapter. New York, NY, USA: Association for Computing Machinery;

50. Sun J, Lu F, Luo Y, Bie L, Xu L, Wang Y. OrthoVenn3: an integrated platform for exploring and visualizing orthologous data across genomes. Nucleic Acids Res. Oxford Academic; 2023; doi: 10.1093/nar/gkad313.

51. Emms DM, Kelly S. OrthoFinder: phylogenetic orthology inference for comparative genomics. Genome Biology. 2019; doi: 10.1186/s13059-019-1832-y.

52. Buchfink B, Reuter K, Drost H-G. Sensitive protein alignments at tree-of-life scale using DIAMOND. Nat Methods. Nature Publishing Group; 2021; doi: 10.1038/s41592-021-01101-x.

53. Mendes FK, Vanderpool D, Fulton B, Hahn MW. CAFE 5 models variation in evolutionary rates among gene families. Bioinformatics. 2021; doi: 10.1093/bioinformatics/btaa1022.

54. Paterson AH, Wendel JF, Gundlach H, Guo H, Jenkins J, Jin D, et al.. Repeated polyploidization of Gossypium genomes and the evolution of spinnable cotton fibres. Nature. 2012; doi: 10.1038/nature11798.

55. Cheng C-Y, Krishnakumar V, Chan AP, Thibaud-Nissen F, Schobel S, Town CD. Araport11: a complete reannotation of the Arabidopsis thaliana reference genome. Plant J. 2017; doi: 10.1111/tpj.13415.

56. Klopfenstein DV, Zhang L, Pedersen BS, Ramírez F, Vesztrocy AW, Naldi A, et al.. GOATOOLS: A Python library for Gene Ontology analyses. Sci Rep. 2018; doi: 10.1038/s41598-018-28948-z.

57. Colonges K, Loor Solorzano RG, Jimenez J-C, Lahon M-C, Seguine E, Calderon D, et al.. Variability and genetic determinants of cocoa aromas in trees native to South Ecuadorian Amazonia. *PLANTS, PEOPLE*, PLANET. 2022; doi: 10.1002/ppp3.10268.

58. Binns D, Dimmer E, Huntley R, Barrell D, O’Donovan C, Apweiler R. QuickGO: a web-based tool for Gene Ontology searching. Bioinformatics. 2009; doi: 10.1093/bioinformatics/btp536.

59. Hardie DG. PLANT PROTEIN SERINE/THREONINE KINASES: Classification and Functions. Annu Rev Plant Physiol Plant Mol Biol. 1999; doi: 10.1146/annurev.arplant.50.1.97.

60. Dantas LG, Guerra M. Chromatin differentiation between Theobroma cacao L. and T. grandiflorum Schum. Genet Mol Biol. 2010; doi: 10.1590/S1415-47572009005000103.

61. Jedlicka P, Lexa M, Kejnovsky E. What Can Long Terminal Repeats Tell Us About the Age of LTR Retrotransposons, Gene Conversion and Ectopic Recombination? Frontiers in Plant Science. 112020;

62. Pedro DLF, Amorim TS, Varani A, Guyot R, Domingues DS, Paschoal AR. An Atlas of Plant Transposable Elements. F1000Res. 2021; doi: 10.12688/f1000research.74524.1.

63. Bourque G, Burns KH, Gehring M, Gorbunova V, Seluanov A, Hammell M, et al.. Ten things you should know about transposable elements. Genome Biology. 2018; doi: 10.1186/s13059-018-1577-z.

64. Lanaud C, Fouet O, Legavre T, Lopes U, Sounigo O, Eyango MC, et al.. Deciphering the Theobroma cacao self-incompatibility system: from genomics to diagnostic markers for self-compatibility. Journal of Experimental Botany. 2017; doi: 10.1093/jxb/erx293.

65. Stirnimann CU, Petsalaki E, Russell RB, Müller CW. WD40 proteins propel cellular networks. Trends Biochem Sci. 2010; doi: 10.1016/j.tibs.2010.04.003.

66. Alandete-Saez M, Ron M, Leiboff S, McCormick S. Arabidopsis thaliana GEX1 has dual functions in gametophyte development and early embryogenesis. Plant J. 2011; doi: 10.1111/j.1365-313X.2011.04713.x.

67. Alvim P de T. CHAPTER 10 - Cacao. In: Alvim P de T, Kozlowski TT, editors. Ecophysiology of Tropical Crops. Academic Press;

68. Romero Vergel AP, Camargo Rodriguez AV, Ramirez OD, Arenas Velilla PA, Gallego AM. A Crop Modelling Strategy to Improve Cacao Quality and Productivity. Plants (Basel). 2022; doi: 10.3390/plants11020157.

69. Nicolau M, Picault N, Descombin J, Jami-Alahmadi Y, Feng S, Bucher E, et al.. The plant mobile domain proteins MAIN and MAIL1 interact with the phosphatase PP7L to regulate gene expression and silence transposable elements in Arabidopsis thaliana. PLoS Genet. 2020; doi: 10.1371/journal.pgen.1008324.

70. G Ea, A Wm, P Em, Rojano BA, A Jjm. Caracterización y extracción lipídica de las semillas del cacao amazónico [Theobroma grandiflorum]. Ciencia en Desarrollo. 2016; doi: 10.19053/01217488.4237.

71. Lam KC, Ibrahim RK, Behdad B, Dayanandan S. Structure, function, and evolution of plant O-methyltransferases. Genome. 2007; doi: 10.1139/g07-077.

72. Guillaumie S, Ilg A, Réty S, Brette M, Trossat-Magnin C, Decroocq S, et al.. Genetic analysis of the biosynthesis of 2-methoxy-3-isobutylpyrazine, a major grape-derived aroma compound impacting wine quality. Plant Physiol. 2013; doi: 10.1104/pp.113.218313.

73. Mathiazhagan M, Chidambara B, Hunashikatti LR, Ravishankar KV. Genomic Approaches for Improvement of Tropical Fruits: Fruit Quality, Shelf Life and Nutrient Content. Genes (Basel). 2021; doi: 10.3390/genes12121881.

74. Wu B, Liu X, Xu K, Zhang B. Genome-wide characterization, evolution and expression profiling of UDP-glycosyltransferase family in pomelo (Citrus grandis) fruit. BMC Plant Biol. 2020; doi: 10.1186/s12870-020-02655-2.

75. Mendez-Yañez A, Ramos P, Morales-Quintana L. Role of Glycoproteins during Fruit Ripening and Seed Development. Cells. 2021; doi: 10.3390/cells10082095.

76. Bilal Tufail M, Yasir M, Zuo D, Cheng H, Ali M, Hafeez A, et al.. Identification and Characterization of Phytocyanin Family Genes in Cotton Genomes. Genes (Basel). 2023; doi: 10.3390/genes14030611.

77. Hashiguchi T, Sakakibara Y, Hara Y, Shimohira T, Kurogi K, Akashi R, et al.. Identification and characterization of a novel kaempferol sulfotransferase from Arabidopsis thaliana. Biochem Biophys Res Commun. 2013; doi: 10.1016/j.bbrc.2013.04.022.

78. Melo CWB de, Bandeira M de J, Maciel LF, Bispo E da S, Souza CO de, Soares SE. Chemical composition and fatty acids profile of chocolates produced with different cocoa (*Theobroma cacao L*.) cultivars. Food Sci Technol. Sociedade Brasileira de Ciência e Tecnologia de Alimentos; 2020; doi: 10.1590/fst.43018.

79. Cohen K de O, Jackix M de NH. Características químicas e física da gordura de cupuaçu e da manteiga de cacau. Planaltina, DF: Embrapa Cerrados, 2009.; 2009;

80. Mostafa S, Wang Y, Zeng W, Jin B. Floral Scents and Fruit Aromas: Functions, Compositions, Biosynthesis, and Regulation. Frontiers in Plant Science. 132022;

81. Forlani S, Masiero S, Mizzotti C. Fruit ripening: the role of hormones, cell wall modifications, and their relationship with pathogens. Journal of Experimental Botany. 2019; doi: 10.1093/jxb/erz112.

82. Qiao X, Zhang S, Paterson AH. Pervasive genome duplications across the plant tree of life and their links to major evolutionary innovations and transitions. Computational and Structural Biotechnology Journal. 2022; doi: 10.1016/j.csbj.2022.06.026.

83. Wang J, Tao F, Marowsky NC, Fan C. Evolutionary Fates and Dynamic Functionalization of Young Duplicate Genes in Arabidopsis Genomes. Plant Physiol. 2016; doi: 10.1104/pp.16.01177.

84. Kono TJY, Brohammer AB, McGaugh SE, Hirsch CN. Tandem Duplicate Genes in Maize Are Abundant and Date to Two Distinct Periods of Time. G3 (Bethesda). 2018; doi: 10.1534/g3.118.200580.

85. Duarte JM, Wall PK, Edger PP, Landherr LL, Ma H, Pires JC, et al.. Identification of shared single copy nuclear genes in Arabidopsis, Populus, Vitis and Oryza and their phylogenetic utility across various taxonomic levels. BMC Evol Biol. 2010; doi: 10.1186/1471-2148-10-61.

86. Panchy N, Lehti-Shiu M, Shiu S-H. Evolution of Gene Duplication in Plants. Plant Physiol. 2016; doi: 10.1104/pp.16.00523.

87. Renny-Byfield S, Rodgers-Melnick E, Ross-Ibarra J. Gene Fractionation and Function in the Ancient Subgenomes of Maize. Mol Biol Evol. 2017; doi: 10.1093/molbev/msx121.

88. Rastogi S, Liberles DA. Subfunctionalization of duplicated genes as a transition state to neofunctionalization. BMC Evol Biol. 2005; doi: 10.1186/1471-2148-5-28.

89. Zamora A, Sun Q, Hamblin MT, Aquadro CF, Kresovich S. Positively selected disease response orthologous gene sets in the cereals identified using Sorghum bicolor L. Moench expression profiles and comparative genomics. Mol Biol Evol. 2009; doi: 10.1093/molbev/msp114.

90. Rech GE, Vargas WA, Sukno SA, Thon MR. Identification of positive selection in disease response genes within members of the Poaceae. Plant Signal Behav. 2012; doi: 10.4161/psb.22362.

91. Yoshikuni Y, Martin VJJ, Ferrin TE, Keasling JD. Engineering cotton (+)-delta-cadinene synthase to an altered function: germacrene D-4-ol synthase. Chem Biol. 2006; doi: 10.1016/j.chembiol.2005.10.016.

92. Karaca M, Ince AG. Grafting-induced seed gossypol levels by demethylation of (+)-delta-cadinene synthase genes in upland cotton. Plant Breeding. 2023; doi: 10.1111/pbr.13066.

93. Colonges K, Jimenez J-C, Saltos A, Seguine E, Loor Solorzano RG, Fouet O, et al.. Two Main Biosynthesis Pathways Involved in the Synthesis of the Floral Aroma of the Nacional Cocoa Variety. Front Plant Sci. 2021; doi: 10.3389/fpls.2021.681979.

94. Zhu B-Q, Cai J, Wang Z-Q, Xu X-Q, Duan C-Q, Pan Q-H. Identification of a plastid-localized bifunctional nerolidol/linalool synthase in relation to linalool biosynthesis in young grape berries. Int J Mol Sci. 2014; doi: 10.3390/ijms151221992.

95. Kiryu M, Hamanaka M, Yoshitomi K, Mochizuki S, Akimitsu K, Gomi K. Rice terpene synthase 18 (OsTPS18) encodes a sesquiterpene synthase that produces an antibacterial (E)- nerolidol against a bacterial pathogen of rice. J Gen Plant Pathol. 2018; doi: 10.1007/s10327-018-0774-7.

96. Salehi B, Fokou PVT, Sharifi-Rad M, Zucca P, Pezzani R, Martins N, et al.. The Therapeutic Potential of Naringenin: A Review of Clinical Trials. Pharmaceuticals (Basel). 2019; doi: 10.3390/ph12010011.

97. Carmona-Hernandez JC, Le M, Idárraga-Mejía AM, González-Correa CH. Flavonoid/Polyphenol Ratio in Mauritia flexuosa and Theobroma grandiflorum as an Indicator of Effective Antioxidant Action. Molecules. 2021; doi: 10.3390/molecules26216431.

98. Liu Y-W, Han C-H, Lee M-H, Hsu F-L, Hou W-C. Patatin, the tuber storage protein of potato (Solanum tuberosum L.), exhibits antioxidant activity in vitro. J Agric Food Chem. 2003; doi: 10.1021/jf030016j.

99. Gambuti A, Rinaldi A, Moio L. Use of patatin, a protein extracted from potato, as alternative to animal proteins in fining of red wine. Eur Food Res Technol. 2012; doi: 10.1007/s00217-012-1791-y.

100. Gelley S, Lankry H, Glusac J, Fishman A. Yeast-derived potato patatins: Biochemical and biophysical characterization. Food Chem. 2022; doi: 10.1016/j.foodchem.2021.130984.

101. Liu W, Zhang R, Xiang C, Zhang R, Wang Q, Wang T, et al.. Transcriptomic and Physiological Analysis Reveal That α-Linolenic Acid Biosynthesis Responds to Early Chilling Tolerance in Pumpkin Rootstock Varieties. Front Plant Sci. 2021; doi: 10.3389/fpls.2021.669565.

102. Teerawanichpan P, Qiu X. Fatty acyl-CoA reductase and wax synthase from Euglena gracilis in the biosynthesis of medium-chain wax esters. Lipids. 2010; doi: 10.1007/s11745-010-3395-2.

103. Li W, Liu J, Zhang H, Liu Z, Wang Y, Xing L, et al.. Plant pan-genomics: recent advances, new challenges, and roads ahead. Journal of Genetics and Genomics. 2022; doi: 10.1016/j.jgg.2022.06.004.

104. Clark JW. Genome evolution in plants and the origins of innovation. New Phytol. 2023; doi: 10.1111/nph.19242.

105. Flagel LE, Wendel JF. Gene duplication and evolutionary novelty in plants. New Phytol. 2009; doi: 10.1111/j.1469-8137.2009.02923.x.

106. Birchler JA, Yang H. The multiple fates of gene duplications: Deletion, hypofunctionalization, subfunctionalization, neofunctionalization, dosage balance constraints, and neutral variation. Plant Cell. 2022; doi: 10.1093/plcell/koac076.

