## Supplementary Information and Materials for "Genomic decoding of *Theobroma grandiflorum* (cupuassu) at chromosomal scale: Evolutionary insights for horticultural innovation"

**Running title:** *Theobroma grandiflorum genome*

### List of Supplementary Information Provided

**Supplementary Information 1.** HMW DNA extraction, Sequencing QC, Bioinformatics procedures used to annotate *Theobroma grandiflorum, T. cacao* and *Herrania umbratica* genomes, and additional notes.

**Figure S1.** Distribution of the ncRNA identified in *Theobroma grandiflorum* genome.

**Figure S2.** LTR insertion time of Gypsy and Copia elements. **A.** *Theobroma grandiflorum,* **B.** *T. cacao, and* **C.** *Herrania umbratica.* The vertical black line represents the median, and the dotted line represents the mean.

**Figure S3.** TE_density analyses of all *Theobroma grandiflorum* chromosomes.

**Figure S4. A.** Microsynteny and coliniarity example of subtelomeric regions of *T. grandiflorum*, *T. cacao* and *H. umbratica*, **B.** Microsynteny and coliniarity example of pericentromeric regions of *T. grandiflorum*, *T. cacao* and *H. umbratica.* Blue represents genes in the forward direction, green indicates genes in the reverse direction, and orange denotes transposable elements (TEs).

**Figure S5.** Alignment of GEX1 gene from CH4 loci generated on Jalview [1].

**Figure S6.** Box-plot and swarmplot showing the the Ka/Ks ratio distributions of the selected GO terms associated with fruit traits and defense mechanisms. **A.** *Theobroma cacao*, **B.** *Herrania umbratica*.

**Figure S7.** Genomic mapping of plant disease resistant genes in *Theobroma grandiflorum* chromosomes. Genes under positive selection are shown in red. The cupuassu WBD-resistant QTL is shown in blue.

**Table S1.** GenBank SRA accession numbers used for transcriptome assembly. **A.** All *Theobroma cacao* RNAseq data used. **B.** *Herrania umbratica* RNAseq data used.

**Table S2.** Genome assembly statistics and completeness scores of the three Malvaceae.

**Table S3.** Genome annotation features and statistics of the three Malvaceae.

**Table S4.** Retrocopies identified in *Theobroma grandiflorum*, *T. cacao*, and *Herrania umbratica,* with associated raw data.

**Table S5.** Genome structural features and Statistics for each *Theobroma grandiflorum* chromosome.

**Table S6.** Tranposable elements summary table and statistics identified in *Theobroma grandiflorum*, *T. cacao*, and *Herrania umbratica.*

**Table S7.** Exclusive gene families identified for each genome analyzed.

**Table S8.** Singletons identified in each genome analyzed.

**Table S9.** Expanded and Contracted gene families identified in each genome analyzed.

**Table S10.** GO enrichment analyses raw data.

**Table S11.** Genes and GO terms identified as positively selected by Ka/Ks analysis.

**Table S12.** Gene content and features of cupuassu WBD-resistant QTL.

### 1. Supplementary Information

#### 1.1. DNA and RNA Sequencing and Quality Check

The High Molecular Weight (HMW) DNA, extracted from fresh leaves using the modified CTAB protocol, yielded a concentration of 174 ng/µL, resulting in a total volume of 200 µL and a total quantity of 34.8 µg. The absorbance ratios at 260/280 and 260/230 were 1.74 and 1.43, respectively. The size of the DNA fragments, analyzed using Contour-clamped Homogeneous Electric Field (CHEF) electrophoresis, along with digest quality control, demonstrated the high quality of the obtained HMW DNA (Figure 1).

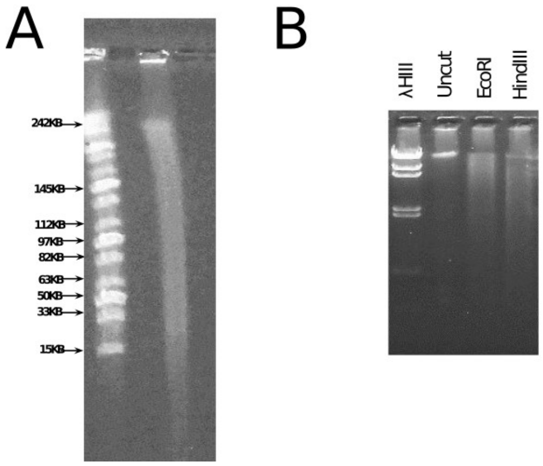


**Figure 1. A.** Size profile on CHEF. **B.** Digest QC.

For the genome assembly, the PacBio Sequel II sequencing produced 1.9 million reads, totaling 30 Gbp, with lengths varying from 43 bp to 48 Kb and an N50 of 15 Kb. The PHRED scores were high, consistently above 50, as illustrated in Figure 2.

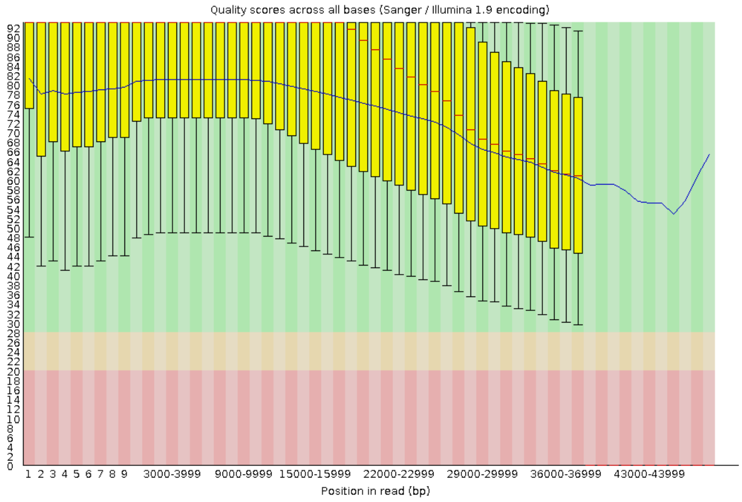


**Figure 2.** FastQC (**https://github.com/s-andrews/FastQC**) report of the DNA sequencing.

For the IsoSeq sequencing, half of a PacBio SMRT cell produced 4.6 million reads, totaling 8.1 Gbp, with lengths ranging from 85 to 12,235 bp and an N50 value of 2 Kb. The mean PHRED score exceeded 40. In contrast, Illumina sequencing yielded 46 million paired-end reads, totaling 9 Gbp, each 2x100 bp, with an average PHRED score greater than 32, as shown in Figure 3 and Figure 4. The obtained RNA concentration was 652 ng/µL, in a total volume of 50 µL, and 32.6 µg of total amount, showinf RNA Integrity Number (RIN) of 7.7.

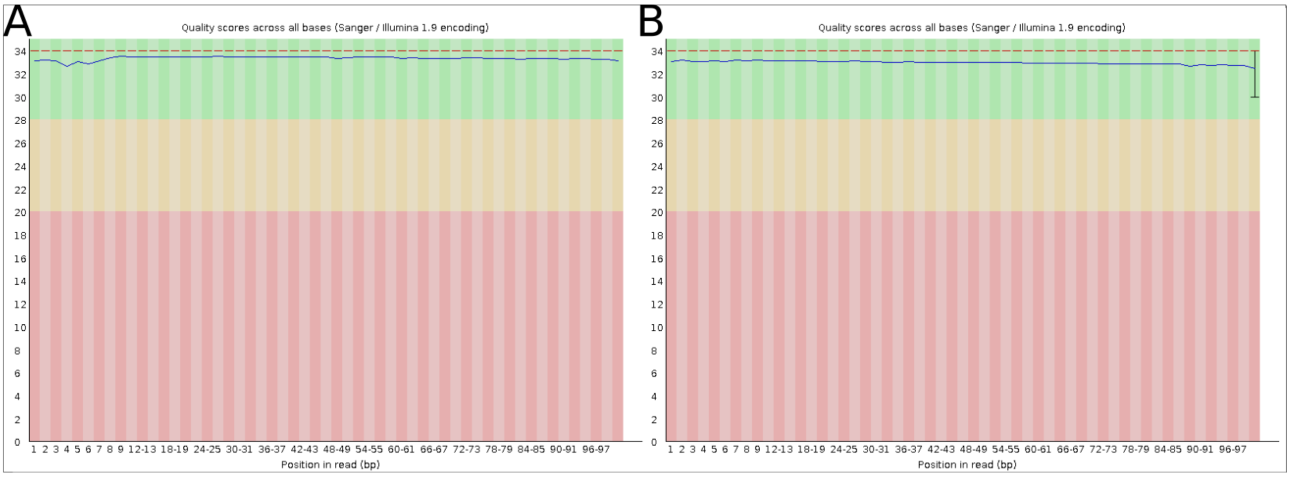


**Figure 3.** FastQC report of the RNAseq Illumina sequencing. **A.** forward reads, **B.** reverse reads.

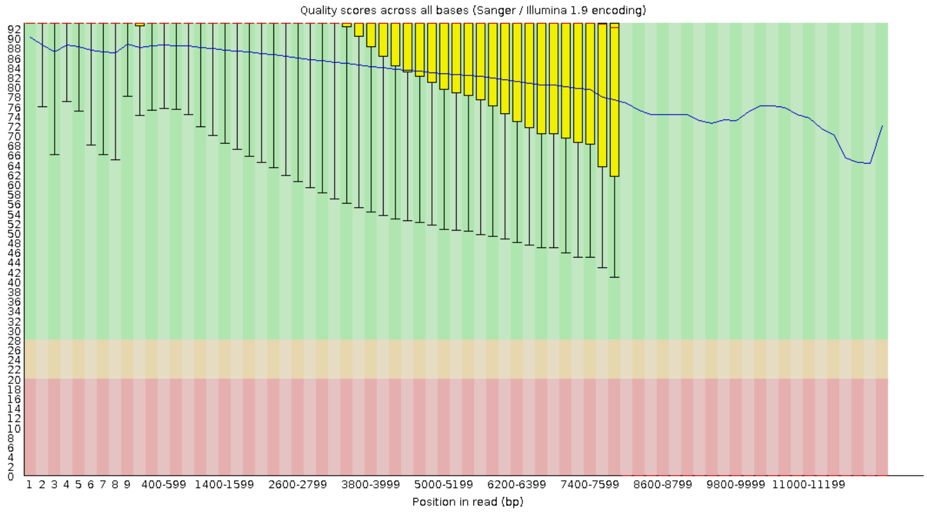


**Figure 4.** FastQC report of the IsoSeq sequencing.

For the Hi-C sequencing, the Illumina NovaSeq platform generated 490 million paired-end reads with a PHRED score exceeding 30.

#### 1.2. Genome Annotation

In the first annotation stage, all Transposable Elements (TEs) were detected and annotated using a modified version of the EDTA pipeline [2] named hereafter as “Plant_Annotation_TEs”, and available and fully documented at following GitHub repository: https://github.com/amvarani/Plant_Annotation_TEs. This modified pipeline is capable to deal with SINE and LINE structural identification using the AnnoSINE (commit: 26301e9) [3] and MGEScan-non-LTR v3.0.0 *[4]*, and also providing autonomous LTR elements full annotation at superfamily and lineages according the nomenclature proposed by Orozco-Arias et al. [5] using TEsorter v1.4.1 [6], and non-automous LTR elements full annotation (*e.g.* LARD, TRIM, TR_GAG, and BARE-2). This pipeline also allows to date the insertion time of each LTR elements and to investigate the evolutionary history of those elements by applying a phylogenetic approach based on IQ-TREE2 inference [7] using maximum likelihood, and generates a soft-masked genome for structural gene annotation.

For structural and functional gene annotation, the second stage generates multiple protein and transcriptome-based evidences using: BRAKER 1+2 [8], BRAKER3 [9], TSEBRA [10], Exonerate v2.4.0 [11], GALBA v1.0.7 [12] + miniprot v0.12 [13], GeMoMa v1.9 [14], all benchmarked by the BUSCO v5.4.5 using the embryophyta_odb10 database [15]. Only evidence showing BUSCO completeness score above 90% (protein mode) were considered for further processing. The approved evidence and transcript structures previously determined using the PASA pipeline were integrated using EvidenceModeler v1.1.0 [16] and further processed in two rounds of the PASA v2.5.3 pipeline to annotate untranslated regions (UTRs), annotation correction, and identification and classification of all identified splicing variations.

A post-processing step was implemented to eliminate false-positive gene predictions that displayed hits to structurally determined TEs as identified in the "Plant_Annotation_TEs" pipeline and potential TE domains according to TEsorter v1.4.1. Moreover, false-positive genes were also removed only if they met all of the following criteria: showed structures shorter than 250 bp; lacked signal peptides as determined by SignalP 6.0 [17]; had no transmembrane domains according to Phobius [18]; showed no alignments with RNA-seq and Iso-Seq data; and exhibited no sequence similarity by BLAST search [19] to entries in the UniProt [20] or NCBI RefSeq [21] plant databases. The BUSCO completeness scores were employed to evaluate the predicted proteins. The gene structure annotation was deemed acceptable only if the score equaled or exceeded the BUSCO genome score.

The approved structural annotation is complemented with the spatial gene arrangement information (*e.g.*, singleton, dispersed, proximal and tandem duplicated, WGD-derived, or transposed genes) generated by MCScanX (commit: b1ca533) [22] and, along with retrocopy identification by RetroScan (commit: cba0f4e) [23] and DupGen_finder (commit: 8001838) [24]. Considering the potential role of retrocopied genes in functional diversification [25], a comparative analysis was conducted, overlapping cupuassu retrocopies with those identified in *T. cacao* and *H. umbratica*

Functional annotation was generated through a BLAST search against public plant databases, such as UniProt, NCBI RefSeq, PlantTFDB v5.0 [26], NLR Atlas [27] and PRGdb v4.0 [28]. EggNOG-mapper v2.1.4-2-4-gb493df4 [29] and InterProScan v5.61-93.0 [30] provide additional functional annotations, including Gene Ontologies (GOs), Enzyme Commission numbers (ECs), Carbohydrate-Active enZYmes (CAZymes), PFAM, and InterPro IDs for each gene and detected isoforms. Metabolic gene clusters are predicted using PlantiSMASH v1.0 [31]. Functional annotation was performed using Blast2GO Basic v6.0 [32]. GOs terms irrelevant to the Viridiplantae clade were subsequently filtered out via the Blast2GO filtering utility. Validation and the generation of the final GFF3 file, encompassing all annotation data, were facilitated by the AGAT (Another Gtf/Gff Analysis Toolkit) package v1.2.0 (https://github.com/NBISweden/AGAT).

The non-coding RNAs (ncRNAs) were annotated using two strategies. The first strategy relied on a BLAST similarity search against the RNAcentral database (version 22) [33]. The second strategy employed a structural search using the Infernal tool [34] in conjunction with the curated RFAM database (version 14.9, 11/2022, encompassing 4,108 families) [35]. Results were filtered based on specific thresholds: low-quality BLAST outcomes were characterized by a query coverage and identity below 95%, while Infernal outcomes were deemed low-quality if they had a bitscore below the covariance model threshold (parameter-cut_ga). The results from both strategies were then integrated to produce the final ncRNA annotation.

The final annotation output includes annotated GFF3 and FASTA nucleotide and protein files, visualized using the JBrowse2 tool [36]. In-house scripts were used to generate the annotation stats. Further details and instructions of the structural and functional annotation pipeline used in this study are provided in figure 5, or in our GitHub repository: https://github.com/amvarani/Plant_Annotation. These same procedures were employed to annotate the *T. cacao* v2 (Criollo cultivar) [37] and *H. umbratica* (Fairchild cultivar) (https://www.ncbi.nlm.nih.gov/datasets/genome/GCF_002168275.1/) genomes.

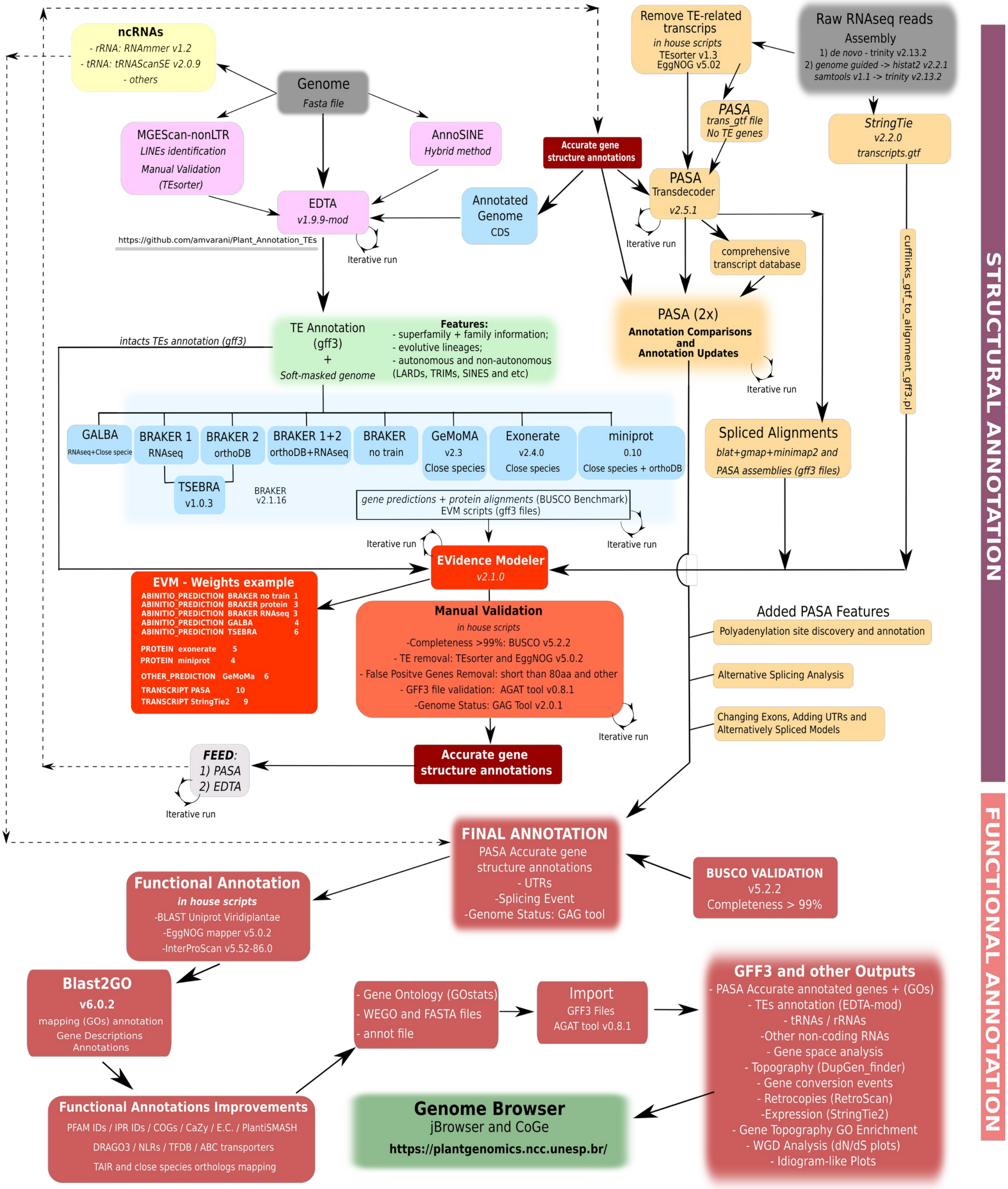


**Figure 5.** Schema of the Plant Genome Annotation Pipeline Employed in This Study. For higher resolution and additional details, please refer to: https://github.com/amvarani/Plant_Annotation.

### 1.3. Whole Genome Duplication and *Theobroma* and *Herrania* species differentiation dating

The core eudicot γ whole-genome triplication event (WGT) is estimated to have occurred approximately 117 million years ago (mya) during the Lower Cretaceous [38]. This event predates the more recent species differentiation, which according to Timetree of Life Database [39] and previous dating studies [40], occurred ~14 mya for *Theobroma* species and ~18 mya between the genera of *Theobroma* and *Herrania*, both during the Miocene epoch.

### 1.4 General Considerations Regarding Comparative Approaches Among *Theobroma grandiflorum*, *T. cacao* v2, and *Herrania umbratica* cultivar Fairchild

The chromosomes and pseudomolecules of *Theobroma cacao* v2 (Belizian Criollo B97-61/B2 cultivar) [37] and *Herrania umbratica* cultivar Fairchild (BioProject: PRJNA383741) are notably shorter than those of *T. grandiflorum*, particularly in the pericentromeric regions (Figure 6). The assembled genomes of *T. cacao* and *H. umbratica* account for only about 70-71% of the estimated genome sizes, as determined by flow cytometry [41,42]. However, previous research indicated minimal variation in chromosome size between *T. cacao* and *T. grandiflorum* [43]. This discrepancy suggests that the genome assemblies of *T. caca*o and *H. umbratica* may be incomplete or unresolved, particularly in complex and highly repetitive regions such as centromeres and pericentromeric areas.

Given that the genomes of *T. cacao* and *H. umbratica* were assembled using earlier sequencing technologies, which are less capable of resolving complex genomic regions, these observations are not surprising. The advancements in long-read sequencing technologies have significantly enhanced our ability to resolve intricate genomic regions. This is evident in the fourth version of the *Musa acuminata* genome [44], and in the resolution of the *T. grandiflorum* genome using HiFi reads and HiC technology.

Therefore, it is imperative to acknowledge that a comparative framework among these three species should consider the various sequencing and assembly strategies utilized. These methodologies may have a direct impact on the structural genomic comparison.

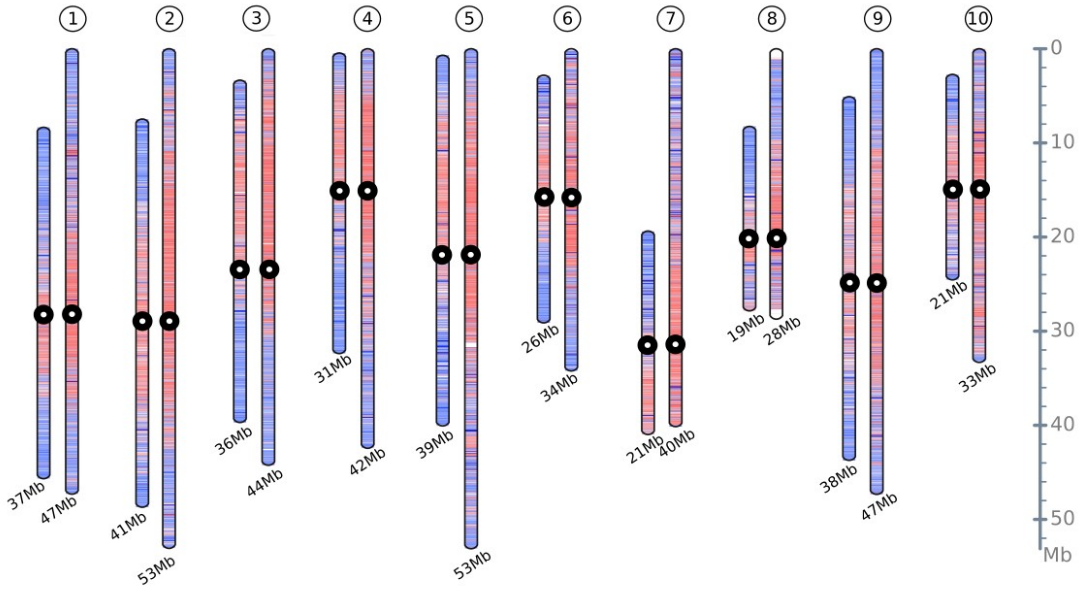


**Figure 6.** Comparative idiogram map between *Theobroma grandiflorum* and *T. cacao* showing the different chromosome sizes. The idiograms illustrate gene-rich regions (blue), TE-rich regions (red), and potential location of centromeres (black circles).

**Figure S1.** Distribution of the ncRNA identified in *Theobroma grandiflorum* genome.

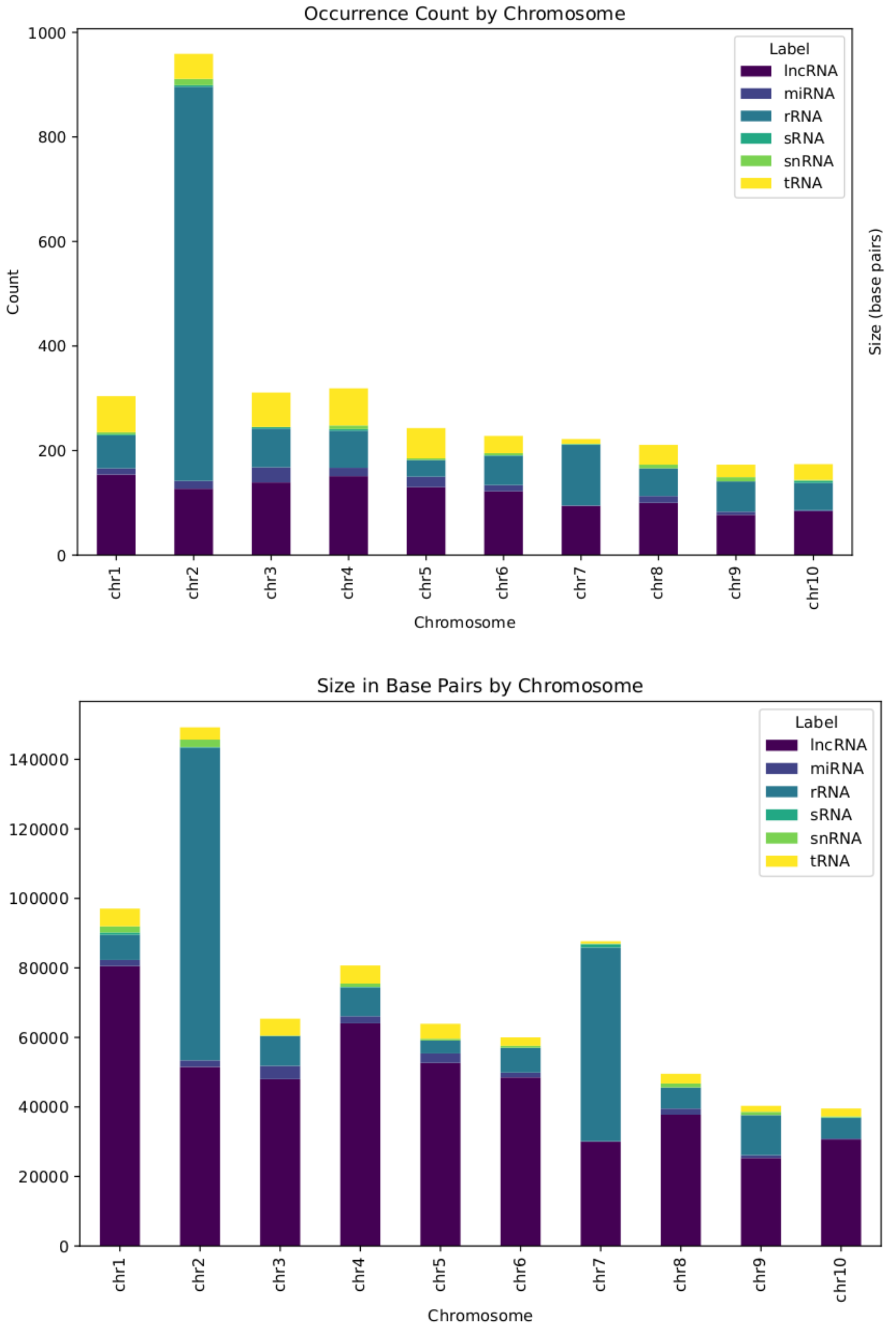

**Figure S2.** LTR insertion time of Gypsy and Copia elements. **A.** *Theobroma grandiflorum,* **B.** *T. cacao, and* **C.** *Herrania umbratica.* The vertical black line represents the median, and the dotted line represents the mean.

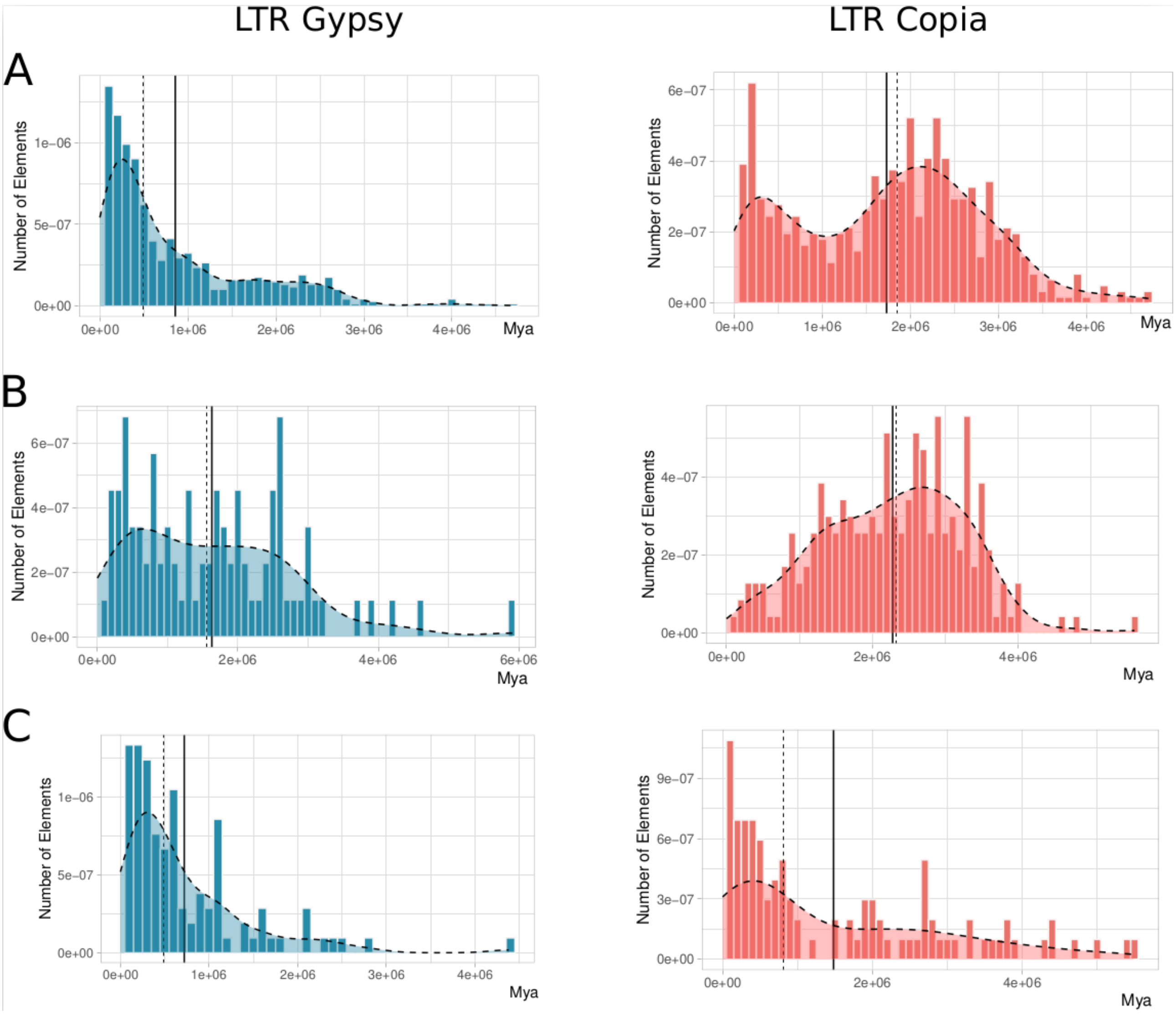

**Figure S3.** TE_density analyses of all *Theobroma grandiflorum* chromosomes.

***Provided as external PDF file***

**Figure S4. A.** Microsynteny and coliniarity example of subtelomeric regions of *T. grandiflorum*, *T. cacao* and *H. umbratica*, **B.** Microsynteny and coliniarity example of pericentromeric regions of *T. grandiflorum*, *T. cacao* and *H. umbratica.* Blue represents genes in the forward direction, green indicates genes in the reverse direction, and orange denotes transposable elements (TEs).

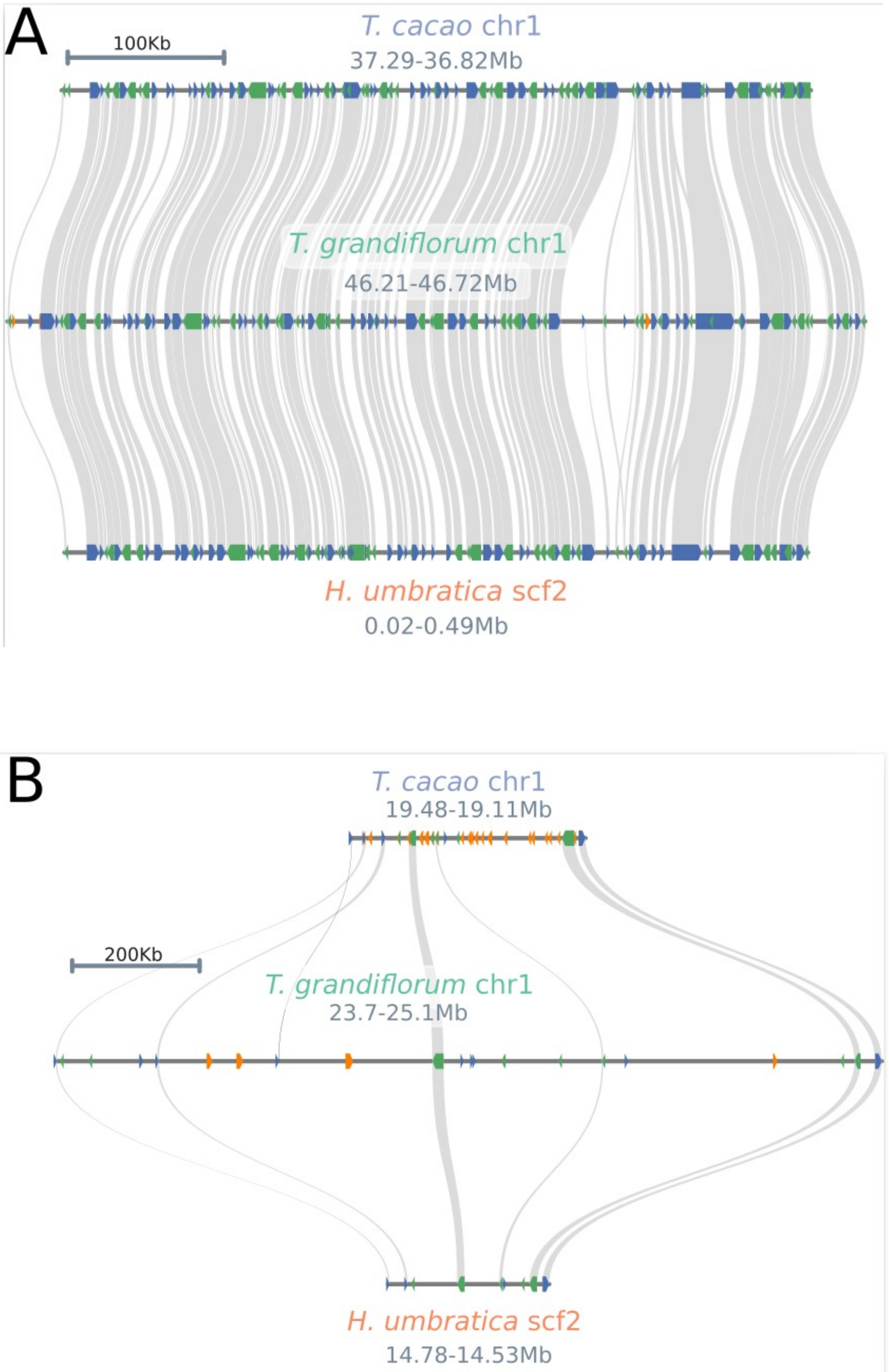

**Figure S5.** Alignment of GEX1 gene from CH4 loci generated on Jalview (Procter et al., 2021).

***Provided as external PDF file***

**Figure S6.** Box-plot and swarmplot showing the the Ka/Ks ratio distributions of the selected GO terms associated with fruit traits and defense mechanisms. A. *Theobroma cacao*, B. *Herrania umbratica*.

***Provided as external PDF file***

**Figure S7.** Genomic mapping of plant disease resistant genes in *Theobroma grandiflorum* chromosomes. Genes under positive selection are shown in red. The cupuassu WBD-resistant QTL is shown in blue.

***Provided as external PDF file***

**Table S1.** GenBank SRA accession numbers used for transcriptome assembly. **A.** All *Theobroma cacao* RNAseq data used. **B.** *Herrania umbratica* RNAseq data used.

**A**

| | | | | | |
|---|---|---|---|---|---|
| SRR7388543 | SRR7388568 | SRR7388593 | SRR11389079 | SRR7172375 | SRR3217284 |
| SRR7388544 | SRR7388569 | SRR7388594 | SRR11389080 | SRR7172376 | SRR3217292 |
| SRR7388545 | SRR7388570 | SRR7388595 | SRR11389081 | SRR7172377 | SRR3217294 |
| SRR7388546 | SRR7388571 | SRR7388596 | SRR11389082 | SRR7172378 | SRR3217297 |
| SRR7388547 | SRR7388572 | SRR7388597 | SRR11389083 | SRR7172379 | SRR3217298 |
| SRR7388548 | SRR7388573 | SRR7388598 | SRR11389084 | SRR7172380 | SRR3217299 |
| SRR7388549 | SRR7388574 | SRR7388599 | SRR7172356 | SRR7172381 | SRR3217301 |
| SRR7388550 | SRR7388575 | SRR7388600 | SRR7172357 | SRR7172382 | SRR3217304 |
| SRR7388551 | SRR7388576 | SRR7388601 | SRR7172358 | SRR7172383 | SRR3217315 |
| SRR7388552 | SRR7388577 | SRR7388602 | SRR7172359 | SRR7172384 | SRR3217317 |
| SRR7388553 | SRR7388578 | SRR7388603 | SRR7172360 | SRR7172385 | SRR3217318 |
| SRR7388554 | SRR7388579 | SRR7388604 | SRR7172361 | SRR7172386 | SRR3217319 |
| SRR7388555 | SRR7388580 | SRR7388605 | SRR7172362 | SRR7172387 | SRR1034656 |
| SRR7388556 | SRR7388581 | SRR7388606 | SRR7172363 | SRR7172388 | SRR1036616 |
| SRR7388557 | SRR7388582 | SRR7388607 | SRR7172364 | SRR7172389 | SRR747762 |
| SRR7388558 | SRR7388583 | SRR7388608 | SRR7172365 | SRR7172390 | SRR747765 |
| SRR7388559 | SRR7388584 | SRR7388609 | SRR7172366 | SRR7172391 | SRR747772 |
| SRR7388560 | SRR7388585 | SRR7388610 | SRR7172367 | SRR3217276 | SRR747773 |
| SRR7388561 | SRR7388586 | SRR7388611 | SRR7172368 | SRR3217277 | SRR747774 |
| SRR7388562 | SRR7388587 | SRR7388612 | SRR7172369 | SRR3217278 | SRR747775 |
| SRR7388563 | SRR7388588 | SRR7388613 | SRR7172370 | SRR3217279 | SRR747776 |
| SRR7388564 | SRR7388589 | SRR7388614 | SRR7172371 | SRR3217280 | SRR747777 |
| SRR7388565 | SRR7388590 | SRR11389076 | SRR7172372 | SRR3217281 | SRR747778 |
| SRR7388566 | SRR7388591 | SRR11389077 | SRR7172373 | SRR3217282 | SRR747779 |
| SRR7388567 | SRR7388592 | SRR11389078 | SRR7172374 | SRR3217283 | SRR13960577 |
| | | | | | SRR13960578 |
| | | | | | SRR13960579 |

**B**

| **SRA Accession** | **Description** |
|---|---|
| SRR5630802 | open flowers |
| SRR5481331 | open flowers |
| SRR5481332 | lateral buds |
| SRR5481333 | closed flower buds |
| SRR5481334 | apical stems |
| SRR5462401 | young leaves |

**Table S2.** Genome assembly statistics and completeness scores of the three Malvaceae.

| Features | *T. grandiflorum* **C1074** (this work) | *T. cacao* **Criollo B97-61/B2 v2** (re-annotation) | *H. umbratica* **Fairchild** (re-assembly and re-annotation) |
|---|---|---|---|
| -Genome length | 423,916,809 bp | 324,761,211 bp | 249,829,655 bp |
| -% assembled by flow cytometry | 94% | 71% | 70% |
| **--- Primary Assembly** | | | |
| -------Assembly size | 434,716,736 | - | 267,828,918 |
| -----Number of contigs | 131 | - | 3464 |
| -----Longest contigs | 53,228,916 | - | 9,298,149 |
| -----Shortest contigs | 17,612 | - | 208 |
| -----Number of contigs > 1K | 131 | - | 3,456 |
| -----Number of contigs > 10K | 131 | - | 1,428 |
| -----Number of contigs > 100K | 38 | - | 320 |
| -----Number of contigs > 1M | 14 | - | 60 |
| -----Number of contigs > 10M | 11 | - | 0 |
| -----Mean contigs size | 33,18,448 | - | 77,318 |
| -----Median contigs size | 46,682 | - | 6,769 |
| -----N50 contigs length | 42,442,329 | - | 1,047,534 |
| -----L50 contigs count | 6 | - | 55 |
| **--- HiC Assembly / GenBank** | | | |
| --- Chromosomes / Scaffolds | 10 | 10 + 1 unplaced | 621 |
| ----- N50/L50 | 44 Mb / 5 | 36 Mb / 5 | 13 Mb / 8 |
| -BUSCO genome | 98.4% | 99.3% | 98.3% |
| -BUSCO annotation | 99.8% | 99.4% | 99.1% |
| -LAI | 15.66 | 9.53 | 14.02 |

**Table S3.** Genome annotation features and statistics of the three Malvaceae.

| **Features** | ***T. grandiflorum* C1074** (this work) | ***T. cacao* Criollo B97-61/B2 v2** (re-annotation) | ***H. umbratica* Fairchild** (re-assembly and re-annotation) |
|---|---|---|---|
| -Number of genes | 31,381 | 30,623 | 27905 |
| -- Number of exons | 285,619 | 271540 | 314599 |
| -- Number of introns | 238,948 | 223339 | 264443 |
| -- Number of CDSs | 46,671 | 48201 | 50156 |
| ---- complete CDS | 46,625 | 46872 | 50049 |
| ---- start, no stop CDS | 8 | 387 | 46 |
| ---- stop, no start CDS | 22 | 453 | 46 |
| ---- no stop, no start CDS | 16 | 489 | 15 |
| -- Overlapping genes | 3233 | 5965 | 4795 |
| -- Contained genes | 892 | 2178 | 1374 |
| -Shortest gene | 153 bp | 150 bp | 153 bp |
| -Longest gene | 227,387 bp | 143,212 bp | 121,875 bp |
| -Longest CDS | 16,371 bp | 16,368 bp | 16,374 bp |
| -mean gene length | 3,374 bp | 3,521 bp | 3,642 bp |
| -mean CDS length | 1,331 bp | 1,214 bp | 1,332 bp |
| -mean exons per gene | 6 | 5 | 6 |
| -mean introns per gene | 5 | 4 | 5 |
| -WGD gene pairs | 4,996 | 4,875 | 8,505 |
| -Tandem gene pairs | 3,667 | 3,476 | 3,146 |
| -Proximal duplicated gene pairs | 3,551 | 2,562 | 1,960 |
| -Dispersed duplicated gene pairs | 14.567 | 14,222 | 9,773 |
| -Singletons | 4,600 | 5,488 | 4,521 |
| -Transposed gene pairs | 2,367 | 2,703 | 1,204 |
| -Retrocopies | 402 | 410 | 309 |
| -- Parental Gene | 214 | 191 | 169 |
| ---- Chimerical | 197 | 215 | 164 |
| ---- Pseudogene | 37 | 40 | 37 |
| ---- Retrogene | 168 | 155 | 108 |
| % of genome covered by genes | 25% | 33.2% | 40.7% |
| % of genome covered by CDS | 14.70% | 18% | 26.7% |
| % of genome covered by TEs | 63.37% | 44.51% | 35.21% |
| % of Class I Elements | 52.84% | 35.41% | 14.24% |
| % of LTR Gypsy | 15.40% | 8.72% | 2.85% |
| % of LTR Copia | 21.43% | 16.78% | 3.77% |
| % of LTR non-autonomous | 14.86% | 9.03% | 6.78% |

| | | | |
|---|---|---|---|
| % of Class II Elements | 2.75% | 3.48% | 3.02% |
| % TIRs | 1.40% | 1.08% | 1.34% |
| % Helitron | 1.36% | 2.39% | 1.67% |

**Table S4.** Retrocopies identified in *Theobroma grandiflorum*, *T. cacao*, and *Herrania umbratica,* with associated raw data.

***Provided as external Excel file***

**Table S5.** Genome structural features and Statistics for each *Theobroma grandiflorum* chromosome.

| ***Theobroma grandiflorum*** Genome Features | chr1 | chr2 | chr3 | chr4 | chr5 | chr6 | chr7 | chr8 | chr9 | chr10 |
|---|---|---|---|---|---|---|---|---|---|---|
| - Synteny against *T.cacao* | 71.05 (35) | 62.02 (40) | 61.15 (42) | 61.42 (39) | 61.24 (40) | 67.57 (37) | 63.18 (38) | 68.95 (37) | 69.38 (36) | 61.76 (38) |
| (% mean (SD) and % median) | 86.67 | 76.92 | 80.00 | 75.00 | 76.92 | 83.33 | 75.00 | 84.62 | 85.71 | 73.33 |
| - Synteny against *H. umbratica* | 66.93 (36) | 53.61 (40) | 57.01 (42) | 58.21 (40) | 56.93 (40) | 61.87 (38) | 53.11 (40) | 62.01 (38) | 65.27 (37) | 57.62 (38) |
| (% mean (SD) and % median) | 81.82 | 62.50 | 71.43 | 71.43 | 67.54 | 75.00 | 63.64 | 75.00 | 80.00 | 66.67 |
| -Genes (count) | 4,199 | 3,764 | 3,258 | 3303 | 3544 | 2550 | 2824 | 2044 | 3624 | 2271 |
| -Genes (% bp occupied) | 24.28 | 17.46 | 19.08 | 20.41 | 18.51 | 21.92 | 19.87 | 20.43 | 21.20 | 18.18 |
| -tRNAs (count) | 74 | 48 | 69 | 75 | 63 | 35 | 9 | 40 | 24 | 34 |
| -snoRNA (count) | 144 | 138 | 120 | 116 | 107 | 92 | 85 | 100 | 92 | 64 |
| -miRNA (count) | 12 | 15 | 29 | 16 | 20 | 12 | 1 | 13 | 6 | 2 |
| -sRNA (count) | 2 | 3 | 3 | 4 | - | 2 | 1 | - | - | 4 |
| -lncRNA (count) | 154 | 127 | 139 | 151 | 130 | 122 | 94 | 100 | 77 | 84 |
| -RNA | | | | | | | | | | |
| --5S (coordinates) | | 21,423,309<br>26,917,293 | | | | | | | | |
| --45S (coordinates) | | | | | | | 4,890<br>504,355 | | | |
| -TEs (count) | 41799 | 52670 | 43272 | 40554 | 56794 | 32476 | 44759 | 25008 | 44134 | 35774 |
| -TEs (% bp occupied) | 48.88 | 59.59 | 57.41 | 56.28 | 60.35 | 55.73 | 61.46 | 57.03 | 51.92 | 59.27 |
| -LTR Copia (count) | 8957 | 12730 | 10116 | 9436 | 12724 | 7505 | 9079 | 5280 | 9732 | 7388 |
| -LTR Copia (% bp occupied) | 16.64 | 21.66 | 20.40 | 19.33 | 20.63 | 19.62 | 18.36 | 16.27 | 17.99 | 18.45 |
| -LTR Gypsy (count) | 7237 | 10507 | 8859 | 7713 | 10609 | 6150 | 7940 | 4420 | 7945 | 6704 |
| -LTR Gypsy (% bp occupied) | 10.89 | 14.39 | 14.47 | 13.27 | 14.19 | 13.55 | 15.09 | 10.77 | 11.99 | 14.90 |
| -LARD (count) | 4818 | 6398 | 5297 | 5015 | 7665 | 3879 | 6344 | 3190 | 4636 | 4631 |
| -LARD (% bp occupied) | 6.04 | 7.42 | 7.01 | 7.36 | 7.63 | 6.94 | 9.60 | 7.15 | 6.09 | 8.44 |
| -TRIM (count) | 227 | 252 | 188 | 240 | 245 | 166 | 220 | 138 | 244 | 202 |
| -TRIM (% bp occupied) | 0.17 | 0.17 | 0.12 | 0.17 | 0.16 | 0.15 | 0.23 | 0.16 | 0.17 | 0.18 |
| -BARE-2 (count) | 572 | 651 | 512 | 539 | 690 | 445 | 450 | 321 | 595 | 352 |
| -BARE-2 (% bp occupied) | 1.00 | 1.12 | 1.02 | 1.19 | 1.07 | 0.93 | 0.93 | 0.82 | 1.02 | 1.06 |
| -TR_GAG (count) | 2110 | 3254 | 2680 | 2376 | 3327 | 1889 | 2356 | 1260 | 2346 | 1966 |
| -TR_GAG (% bp occupied) | 3.62 | 4.81 | 5.07 | 4.73 | 5.25 | 4.38 | 4.48 | 3.42 | 3.74 | 4.51 |
| -LINE (count) | 595 | 708 | 544 | 539 | 873 | 528 | 567 | 377 | 673 | 519 |
| -LINE (% bp occupied) | 0.82 | 0.97 | 0.73 | 0.90 | 1.08 | 0.98 | 0.84 | 0.84 | 0.93 | 1.00 |
| -SINE (count) | 36 | 44 | 31 | 44 | 48 | 22 | 19 | 24 | 36 | 20 |

| | | | | | | | | | | |
|---|---|---|---|---|---|---|---|---|---|---|
| -SINE (% bp occupied) | 0.01 | 0.01 | 0.01 | 0.01 | 0.01 | 0.01 | 0.001 | 0.01 | 0.01 | 0.001 |
| -pararetrovirus (count) | 12 | 36 | 13 | 12 | 30 | 9 | 18 | 34 | 26 | 7 |
| -pararetrovirus (% bp occupied) | 0.02 | 0.10 | 0.06 | 0.04 | 0.08 | 0.03 | 0.08 | 0.18 | 0.09 | 0.06 |
| -TIR (count) | 941 | 929 | 678 | 780 | 877 | 577 | 681 | 498 | 902 | 618 |
| -TIR (% bp occupied) | 1.05 | 1.05 | 0.91 | 1.15 | 0.97 | 0.93 | 1.00 | 1.03 | 1.15 | 1.20 |
| -Helitron (count) | 3074 | 2868 | 2330 | 2333 | 3267 | 2110 | 2174 | 1595 | 3008 | 1863 |
| -Helitron (% bp occupied) | 1.42 | 1.20 | 1.13 | 1.25 | 1.25 | 1.45 | 1.24 | 1.18 | 1.37 | 1.32 |
| -MITE (count) | 812 | 810 | 664 | 721 | 854 | 531 | 692 | 423 | 796 | 573 |
| -MITE (% bp occupied) | 0.30 | 0.26 | 0.26 | 0.26 | 0.34 | 0.25 | 0.26 | 0.25 | 0.29 | 0.28 |
| -Unknown (count) | 12407 | 13483 | 11357 | 10805 | 15582 | 8664 | 14217 | 7448 | 13195 | 10930 |
| -Unknown (% bp occupied) | 6.92 | 6.42 | 6.23 | 6.63 | 7.71 | 6.52 | 9.36 | 14.95 | 7.09 | 7.88 |

**Table S6.** Tranposable elements summary table and statistics identified in *Theobroma grandiflorum*, *T. cacao*, and *Herrania umbratica.*

***Provided as external Excel file***

**Table S7.** Exclusive gene families identified for each genome analyzed.

***Provided as external Excel file***

**Table S8.** Singletons identified in each genome analyzed.

***Provided as external Excel file***

**Table S9.** Expanded and Contracted gene families identified in each genome analyzed.

***Provided as external Excel file***

**Table S10.** GO enrichment analyses raw data.

***Provided as external Excel file***

**Table S11.** Genes and GO terms identified as positively selected by Ka/Ks analysis.

***Provided as external Excel file***

**Table S12.** Gene content and features of cupuassu WBD-resistant QTL.

***Provided as external Excel file***

### References


1. Procter JB, Carstairs GM, Soares B, Mourão K, Ofoegbu TC, Barton D, et al.. Alignment of Biological Sequences with Jalview. *Methods Mol Biol*. 2021; doi: 10.1007/978-1-0716-1036-7_13.

2. Ou S, Su W, Liao Y, Chougule K, Agda JRA, Hellinga AJ, et al.. Benchmarking transposable element annotation methods for creation of a streamlined, comprehensive pipeline. *Genome Biol*. 2019; doi: 10.1186/s13059-019-1905-y.

3. Li Y, Jiang N, Sun Y. AnnoSINE: a short interspersed nuclear elements annotation tool for plant genomes. *Plant Physiol*. 2022; doi: 10.1093/plphys/kiab524.

4. Rho M, Tang H. MGEScan-non-LTR: computational identification and classification of autonomous non-LTR retrotransposons in eukaryotic genomes. *Nucleic Acids Res*. 2009; doi: 10.1093/nar/gkp752.

5. Orozco-Arias S, Isaza G, Guyot R. Retrotransposons in Plant Genomes: Structure, Identification, and Classification through Bioinformatics and Machine Learning. *Int J Mol Sci*. 2019; doi: 10.3390/ijms20153837.

6. Zhang R-G, Li G-Y, Wang X-L, Dainat J, Wang Z-X, Ou S, et al.. TEsorter: an accurate and fast method to classify LTR-retrotransposons in plant genomes. *Hortic Res*. 2022; doi: 10.1093/hr/uhac017.

7. Minh BQ, Schmidt HA, Chernomor O, Schrempf D, Woodhams MD, von Haeseler A, et al.. IQ-TREE 2: New Models and Efficient Methods for Phylogenetic Inference in the Genomic Era. *Mol Biol Evol*. 2020; doi: 10.1093/molbev/msaa015.

8. Hoff KJ, Lomsadze A, Borodovsky M, Stanke M. Whole-Genome Annotation with BRAKER. *Methods Mol Biol*. 2019; doi: 10.1007/978-1-4939-9173-0_5.

9. Gabriel L, Brůna T, Hoff KJ, Ebel M, Lomsadze A, Borodovsky M, et al.. BRAKER3: Fully automated genome annotation using RNA-Seq and protein evidence with GeneMark-ETP, AUGUSTUS and TSEBRA. *bioRxiv*. 2023; doi: 10.1101/2023.06.10.544449.

10. Gabriel L, Hoff KJ, Brůna T, Borodovsky M, Stanke M. TSEBRA: transcript selector for BRAKER. *BMC Bioinformatics*. 2021; doi: 10.1186/s12859-021-04482-0.

11. Slater GSC, Birney E. Automated generation of heuristics for biological sequence comparison. *BMC Bioinformatics*. 2005; doi: 10.1186/1471-2105-6-31.

12. Brůna T, Li H, Guhlin J, Honsel D, Herbold S, Stanke M, et al.. Galba: genome annotation with miniprot and AUGUSTUS. *BMC Bioinformatics*. 2023; doi: 10.1186/s12859-023-05449-z.

13. Li H. Protein-to-genome alignment with miniprot. *Bioinformatics*. 2023; doi: 10.1093/bioinformatics/btad014.

14. Keilwagen J, Hartung F, Grau J. GeMoMa: Homology-Based Gene Prediction Utilizing Intron Position Conservation and RNA-seq Data. *Methods Mol Biol*. 2019; doi: 10.1007/978-1-4939-9173-0_9.

15. Manni M, Berkeley MR, Seppey M, Zdobnov EM. BUSCO: Assessing Genomic Data Quality and Beyond. *Curr Protoc*. 2021; doi: 10.1002/cpz1.323.

16. Haas BJ, Salzberg SL, Zhu W, Pertea M, Allen JE, Orvis J, et al.. Automated eukaryotic gene structure annotation using EVidenceModeler and the Program to Assemble Spliced Alignments. *Genome Biology*. 2008; doi: 10.1186/gb-2008-9-1-r7.

17. Teufel F, Almagro Armenteros JJ, Johansen AR, Gíslason MH, Pihl SI, Tsirigos KD, et al.. SignalP 6.0 predicts all five types of signal peptides using protein language models. *Nat Biotechnol*. 2022; doi: 10.1038/s41587-021-01156-3.

18. Käll L, Krogh A, Sonnhammer ELL. A combined transmembrane topology and signal peptide prediction method. *J Mol Biol*. 2004; doi: 10.1016/j.jmb.2004.03.016.

19. Camacho C, Coulouris G, Avagyan V, Ma N, Papadopoulos J, Bealer K, et al.. BLAST+: architecture and applications. *BMC Bioinformatics*. 2009; doi: 10.1186/1471-2105-10-421.

20. UniProt Consortium. UniProt: the Universal Protein Knowledgebase in 2023. *Nucleic Acids Res*. 2023; doi: 10.1093/nar/gkac1052.

21. O'Leary NA, Wright MW, Brister JR, Ciufo S, Haddad D, McVeigh R, et al.. Reference sequence (RefSeq) database at NCBI: current status, taxonomic expansion, and functional annotation. *Nucleic Acids Res*. 2016; doi: 10.1093/nar/gkv1189.

22. Wang Y, Tang H, Debarry JD, Tan X, Li J, Wang X, et al.. MCScanX: a toolkit for detection and evolutionary analysis of gene synteny and collinearity. *Nucleic Acids Res*. 2012; doi: 10.1093/nar/gkr1293.

23. Wei Z, Sun J, Li Q, Yao T, Zeng H, Wang Y. RetroScan: An Easy-to-Use Pipeline for Retrocopy Annotation and Visualization. *Frontiers in Genetics*. 122021;

24. Qiao X, Li Q, Yin H, Qi K, Li L, Wang R, et al.. Gene duplication and evolution in recurring polyploidization–diploidization cycles in plants. *Genome Biology*. 2019; doi: 10.1186/s13059-019-1650-2.

25. Flagel LE, Wendel JF. Gene duplication and evolutionary novelty in plants. *New Phytol*. 2009; doi: 10.1111/j.1469-8137.2009.02923.x.

26. Jin J, Tian F, Yang D-C, Meng Y-Q, Kong L, Luo J, et al.. PlantTFDB 4.0: toward a central hub for transcription factors and regulatory interactions in plants. *Nucleic Acids Res*. 2017; doi: 10.1093/nar/gkw982.

27. Martin EC, Ion CF, Ifrimescu F, Spiridon L, Bakker J, Goverse A, et al.. NLRscape: an atlas of plant NLR proteins. *Nucleic Acids Res*. 2023; doi: 10.1093/nar/gkac1014.

28. Calle García J, Guadagno A, Paytuvi-Gallart A, Saera-Vila A, Amoroso CG, D'Esposito D, et al.. PRGdb 4.0: an updated database dedicated to genes involved in plant disease resistance process. *Nucleic Acids Res*. 2022; doi: 10.1093/nar/gkab1087.

29. Huerta-Cepas J, Szklarczyk D, Heller D, Hernández-Plaza A, Forslund SK, Cook H, et al.. eggNOG 5.0: a hierarchical, functionally and phylogenetically annotated orthology resource based on 5090 organisms and 2502 viruses. *Nucleic Acids Res*. 2019; doi: 10.1093/nar/gky1085.

30. Mulder N, Apweiler R. InterPro and InterProScan: tools for protein sequence classification and comparison. *Methods Mol Biol*. 396:59–702007;

31. Kautsar SA, Suarez Duran HG, Medema MH. Genomic Identification and Analysis of Specialized Metabolite Biosynthetic Gene Clusters in Plants Using PlantiSMASH. *Methods Mol Biol*. 2018; doi: 10.1007/978-1-4939-7874-8_15.

32. Conesa A, Götz S, García-Gómez JM, Terol J, Talón M, Robles M. Blast2GO: a universal tool for annotation, visualization and analysis in functional genomics research. *Bioinformatics*. 2005; doi: 10.1093/bioinformatics/bti610.

33. RNAcentral Consortium. RNAcentral 2021: secondary structure integration, improved sequence search and new member databases. *Nucleic Acids Res*. 2021; doi: 10.1093/nar/gkaa921.

34. Nawrocki EP, Eddy SR. Infernal 1.1: 100-fold faster RNA homology searches. *Bioinformatics*. 2013; doi: 10.1093/bioinformatics/btt509.

35. Kalvari I, Nawrocki EP, Ontiveros-Palacios N, Argasinska J, Lamkiewicz K, Marz M, et al.. Rfam 14: expanded coverage of metagenomic, viral and microRNA families. *Nucleic Acids Res*. 2021; doi: 10.1093/nar/gkaa1047.

36. Diesh C, Stevens GJ, Xie P, De Jesus Martinez T, Hershberg EA, Leung A, et al.. JBrowse 2: a modular genome browser with views of synteny and structural variation. *Genome Biol*. 2023; doi: 10.1186/s13059-023-02914-z.

37. Argout X, Martin G, Droc G, Fouet O, Labadie K, Rivals E, et al.. The cacao Criollo genome v2.0: an improved version of the genome for genetic and functional genomic studies. *BMC Genomics*. 2017; doi: 10.1186/s12864-017-4120-9.

38. Jiao Y, Leebens-Mack J, Ayyampalayam S, Bowers JE, McKain MR, McNeal J, et al.. A genome triplication associated with early diversification of the core eudicots. *Genome Biol*. 2012; doi: 10.1186/gb-2012-13-1-r3.

39. Kumar S, Suleski M, Craig JM, Kasprowicz AE, Sanderford M, Li M, et al.. TimeTree 5: An Expanded Resource for Species Divergence Times. *Molecular Biology and Evolution*. 2022; doi: 10.1093/molbev/msac174.

40. Richardson JE, Whitlock BA, Meerow AW, Madriñán S. The age of chocolate: a diversification history of Theobroma and Malvaceae. *Frontiers in Ecology and Evolution*. 32015;

41. Argout X, Salse J, Aury J-M, Guiltinan MJ, Droc G, Gouzy J, et al.. The genome of Theobroma cacao. *Nat Genet*. 2011; doi: 10.1038/ng.736.

42. da Silva RA, Souza G, Lemos LSL, Lopes UV, Patrocínio NGRB, Alves RM, et al.. Genome size, cytogenetic data and transferability of EST-SSRs markers in wild and cultivated species of the genus Theobroma L. (Byttnerioideae, Malvaceae). *PLoS One*. 2017; doi: 10.1371/journal.pone.0170799.

43. Dantas LG, Guerra M. Chromatin differentiation between Theobroma cacao L. and T. grandiflorum Schum. *Genet Mol Biol*. 2010; doi: 10.1590/S1415-47572009005000103.

44. Belser C, Baurens F-C, Noel B, Martin G, Cruaud C, Istace B, et al.. Telomere-to-telomere gapless chromosomes of banana using nanopore sequencing. *Commun Biol*. 2021; doi: 10.1038/s42003-021-02559-3.
