## Supplementary figures and images for "Genomic decoding of *Theobroma grandiflorum* (cupuassu) at chromosomal scale: Evolutionary insights for horticultural innovation"

### FigureS3

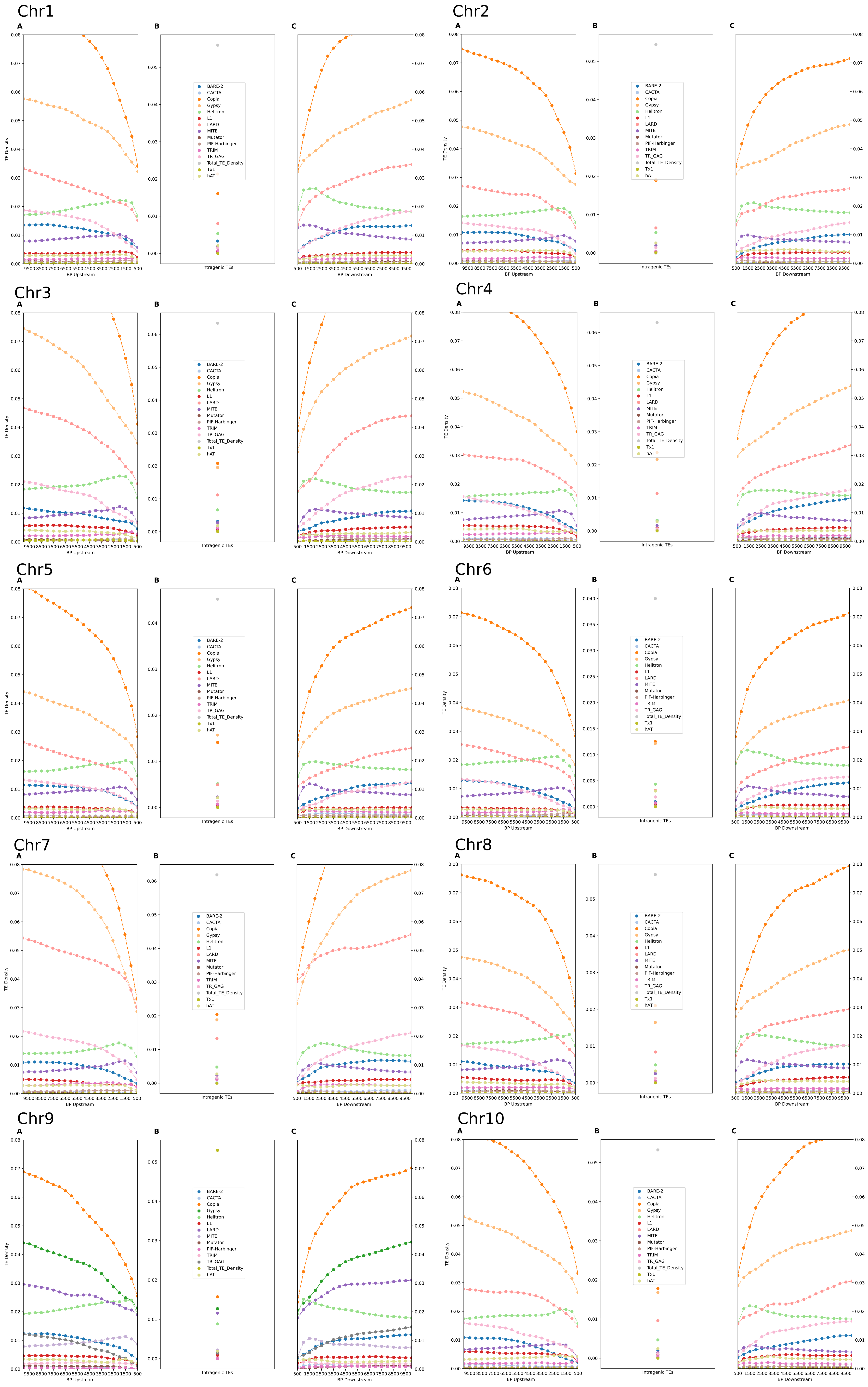

### FigureS5

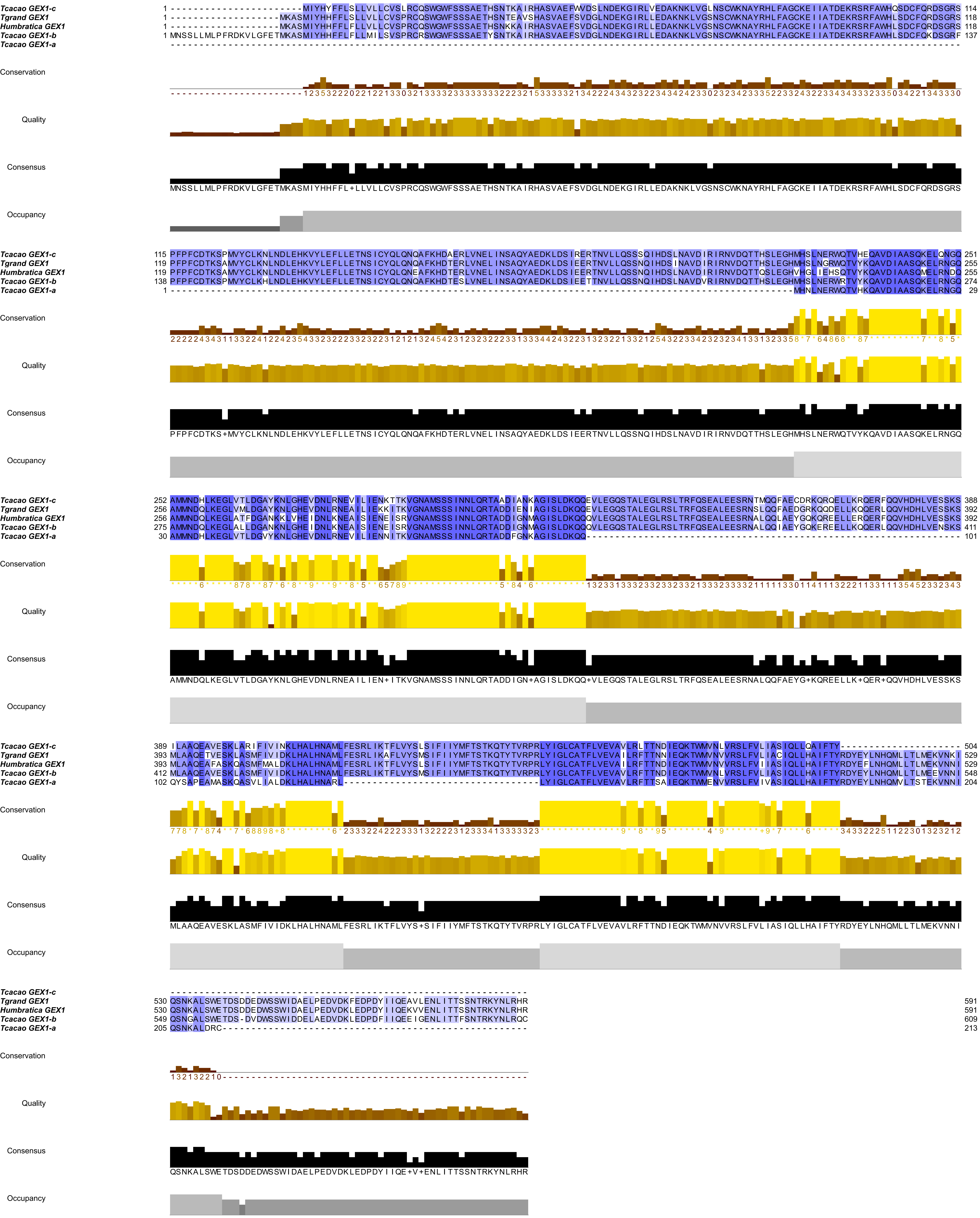

### FigureS6

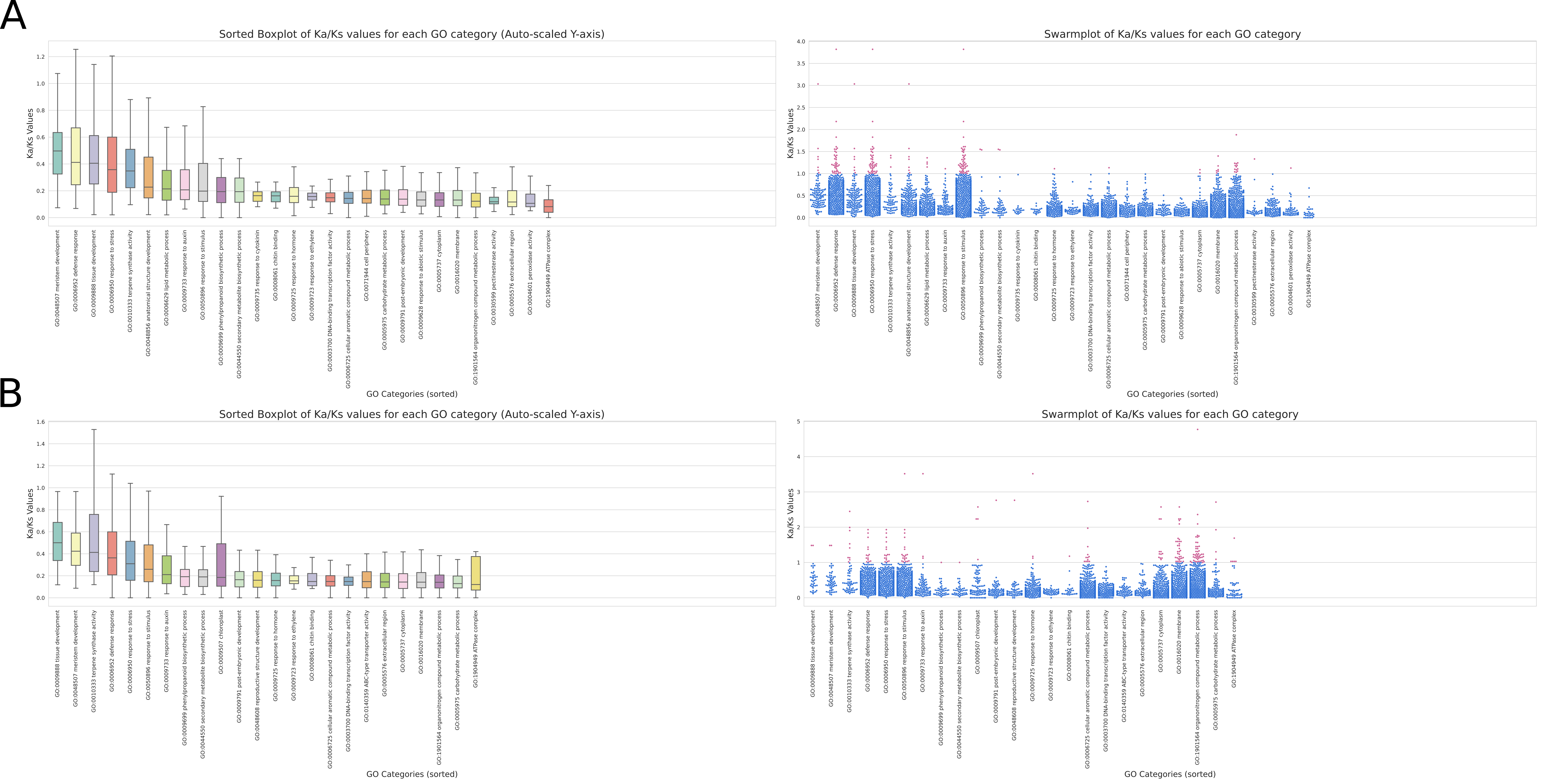

### FigureS7

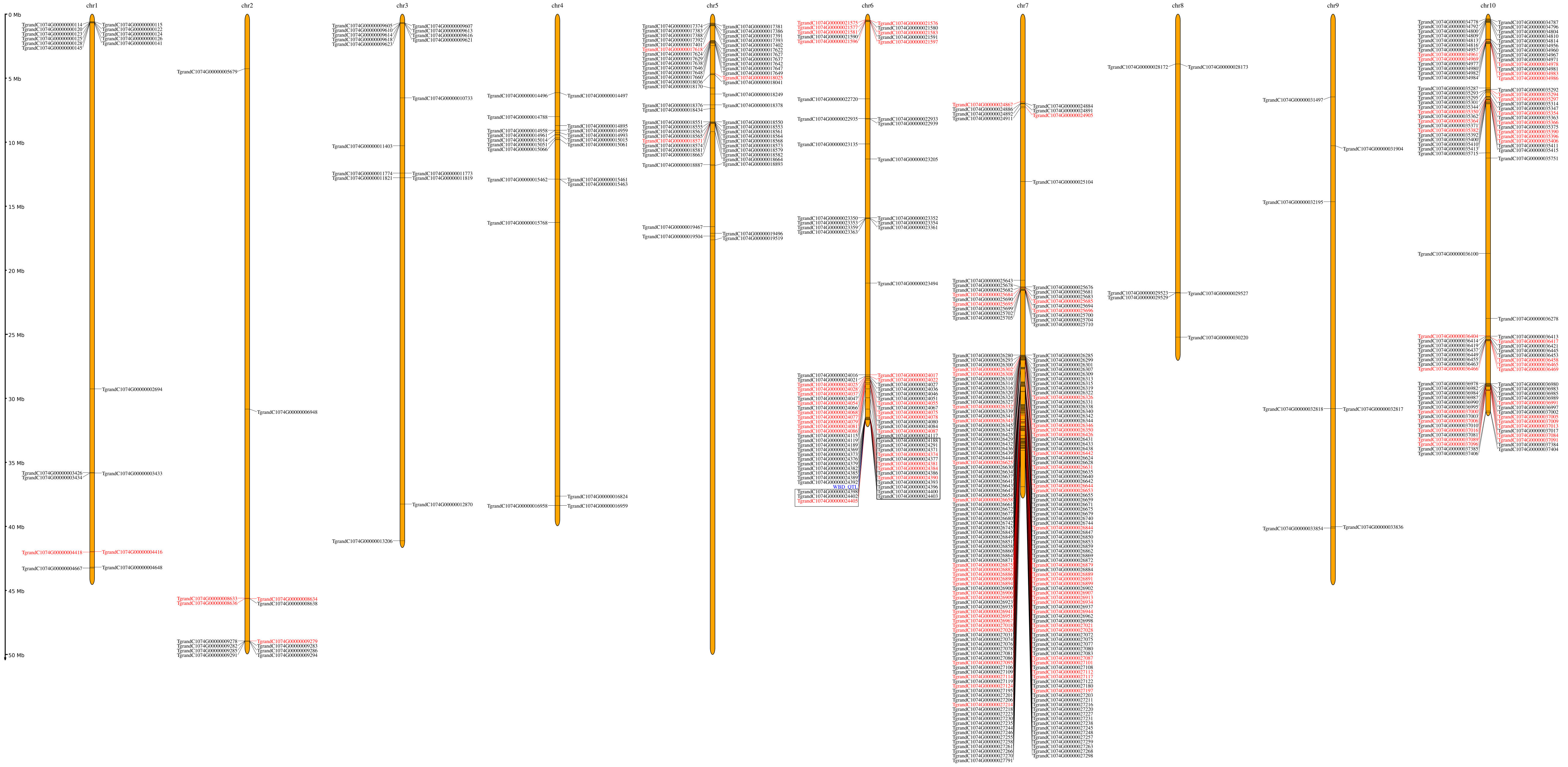
